# The *Drosophila* RNA binding protein Hrp48 binds a specific RNA sequence of the *msl-2* mRNA 3’ UTR to regulate translation

**DOI:** 10.1101/2024.03.26.586676

**Authors:** Andrea Lomoschitz, Julia Meyer, Tanit Guitart, Miroslav Krepl, Karine Lapouge, Clara Hayn, Kristian Schweimer, Bernd Simon, Jiří Šponer, Fátima Gebauer, Janosch Hennig

**Affiliations:** Structural and Computational Biology Unit, European Molecular Biology Laboratory (EMBL), 69117 Heidelberg, Germany; Department of Biochemistry IV – Biophysical Chemistry, University of Bayreuth, 95447 Bayreuth, Germany; Centre for Genomic Regulation (CRG), The Barcelona Institute of Science and Technology, Dr Aiguader 88, 08003 Barcelona, Spain; Institute of Biophysics of the Czech Academy of Sciences, Kralovopolska 135, 612 65 Brno, Czech Republic; Protein Expression and Purification Core Facility, European Molecular Biology Laboratory Heidelberg, 69117 Heidelberg, Germany; Molecular Biology and Biophysics – University of Connecticut Health Center, Farmington, Connecticut 06030, USA; Regional Centre of Advanced Technologies and Materials, Czech Advanced Technology and Research Institute, Palacký University Olomouc, Olomouc 783 71, Czech Republic; Universitat Pompeu Fabra (UPF), 08003 Barcelona, Spain

**Author notes:** These authors contributed equally (Andrea Lomoschitz, Julia Meyer).

**Keywords:** translation regulation, dosage compensation, Hrp48, RNA recognition motif, RNA binding protein

## Abstract

Repression of *msl-2* mRNA translation is essential for viability of *Drosophila melanogaster* females to prevent hypertranscription of both X chromosomes. This translational control event is coordinated by the female-specific protein Sex-lethal (Sxl) which recruits the RNA binding proteins Unr and Hrp48 to the 3’ untranslated region (UTR) of the *msl-2* transcript and represses translation initiation. The mechanism exerted by Hrp48 during translation repression and its interaction with *msl-2* are not well understood. Here we investigate the RNA binding specificity and affinity of the tandem RNA recognition motifs of Hrp48. Using NMR spectroscopy, molecular dynamics simulations and isothermal titration calorimetry, we identified the exact region of *msl-2* 3’ UTR recognized by Hrp48. Additional biophysical experiments and translation assays give further insights into complex formation of Hrp48, Unr, Sxl and RNA. Our results show that Hrp48 binds independent of Sxl and Unr downstream of the E and F binding sites of Sxl and Unr to *msl-2*.

## Introduction

In sexually reproducing organisms the number of X chromosomes across sexes are often inequal. Without further regulation, this would lead to unbalanced expression of X-linked genes (Disteche, 2012). Dosage compensation mechanisms have evolved that, through regulation of transcription and translation, enable uniform expression levels of X-linked genes (Straub and Becker, 2007). In *Drosophila melanogaster*, transcription of the single male X chromosome is upregulated two-fold to equalize the expression levels to that of both female X chromosomes (Graindorge et al., 2011). This hyper-transcription is achieved by the multi-subunit dosage compensation complex (DCC) (Samata and Akhtar, 2018), consisting of five proteins (Mle, Msl1-3, and Mof), of which Msl2 is the rate-limiting component (Belote and Lucchesi, 1980; Gu et al., 1998; Hilfiker et al., 1997) and one of the two long non-coding RNAs *roX1* and *roX2* (Meller and Rattner, 2002). Hyper-transcription would have lethal consequences in females. Here, a mechanism has evolved, which inhibits the assembly of the DCC by translation repression of *msl-2* mRNA (Kelley et al., 1995). This is orchestrated by the female-specific RNA-binding protein (RBP) Sex-lethal (Sxl) in multiple regulatory steps of gene expression (Bashaw and Baker, 1995; Gebauer et al., 2003; Moschall et al., 2017). First, Sxl regulates the alternative splicing of a 5’ untranslated region (UTR) facultative intron of the *msl*-2-pre-mRNA (Förch et al., 2001; Merendino et al., 1999) followed by nuclear retention of *msl-2* transcripts (Gebauer et al., 1998; Graindorge et al., 2013). Second, transcripts that escape to the cytoplasm are repressed at the level of translation initiation by Sxl binding to uridine-rich motifs in both the 5’ and 3’ UTRs of *msl-2* mRNA (designated as A- to F- sites, Figure 1A-C, (Bashaw and Baker, 1997; Beckmann et al., 2005; Gebauer et al., 1998; Gebauer et al., 1999; Kelley et al., 1997; Medenbach et al., 2011)). At the 3’ UTR, in a pAbp-dependent translational control mechanism (Duncan et al., 2009), Sxl together with the RBPs Heterogeneous nuclear ribonucleoprotein 48 (Hrp48, also known as Hrb27C) and Upstream-of-N-ras (Unr) inhibits ribosomal recruitment to the mRNA (Figure 1A, B, (Abaza et al., 2006; Abaza and Gebauer, 2008; Duncan et al., 2006; Duncan et al., 2009; Grskovic et al., 2003; Szostak et al., 2018)). Sxl binds to the E- and F-site of *msl-2* 3’ UTR cooperatively with Unr which acts as a necessary co-repressor (Figure 1C, (Hennig et al., 2014)). As part of the repression complex, Hrp48 binds to the initiation factor eIF3d via direct contacts (Szostak et al., 2018). Ribosomal pre-initiation complexes that could potentially escape the 3’ UTR translational repression are inhibited by Sxl bound to the B-site at the 5’ UTR which interferes with ribosomal scanning and recognition of the AUG initiation codon (Figure 1A, (Beckmann et al., 2005)).

**Figure 1:**
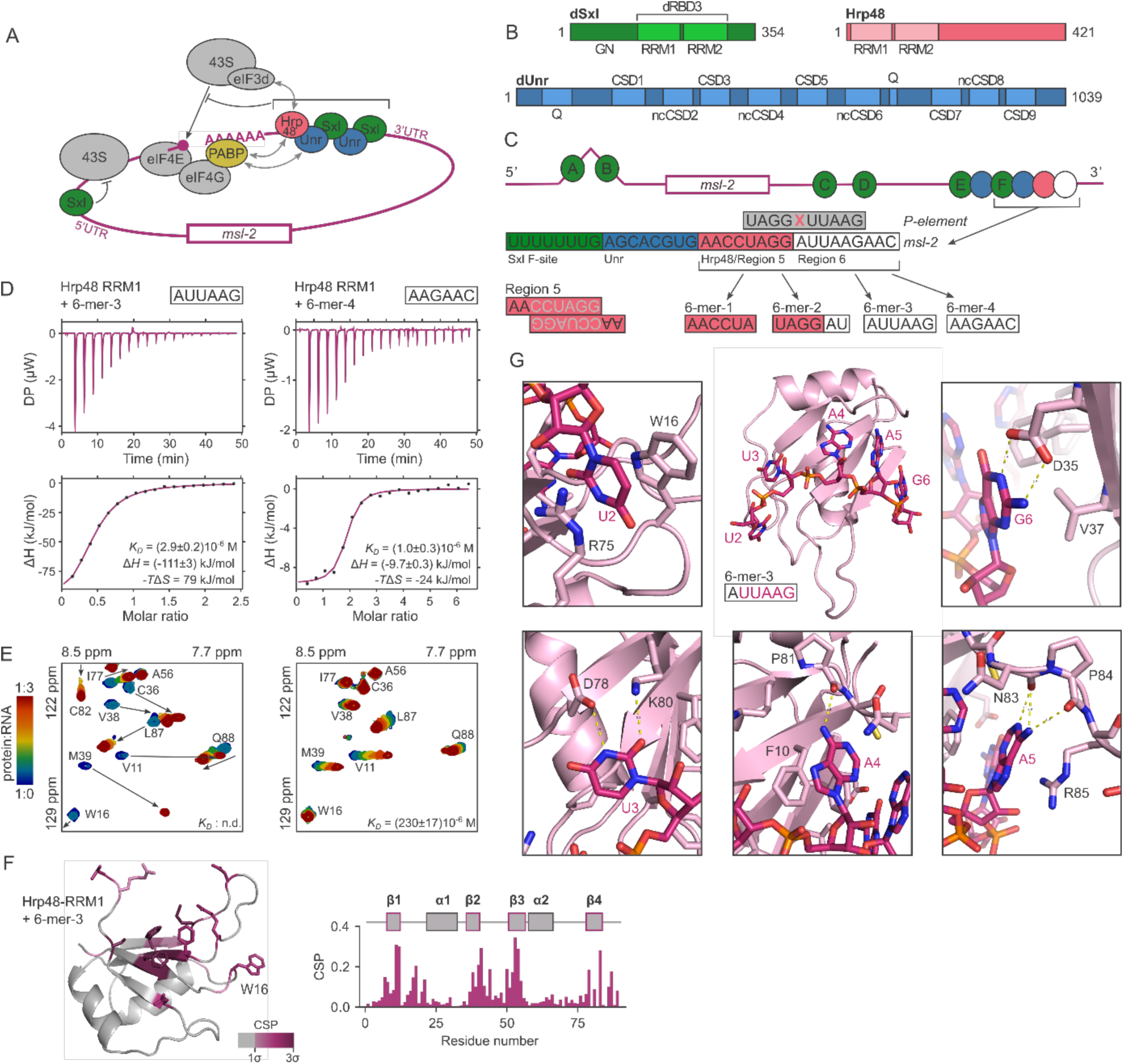
Structure and RNA binding of Hrp48-RRM1. **A:** Model for translation repression of *msl-2* 3’ and 5’ UTR mediated control. Sxl binds to both UTRs and recruits Unr to a specific RNA region of the 3’ UTR, where it inhibits along with Hrp48 the recruitment of the 43S preinitiation complex (PIC). Hrp48 supposedly binds the 43S via interaction with the eIF3d subunit. Those PICs that escaped the 3’ UTR regulation are inhibited by Sxl bound at the 5’ UTR, which prevents scanning of the start codon by the 43S. **B:** Domain arrangement of Sxl, Hrp48 and Unr constructs used in this study. Amino acid numbers are indicated (GN: glycine-asparagine rich region, RRM: RNA recognition motif, dRBD3: former nomenclature of a region encompassing both RRMs of Sxl, residues 123-294, Q: glutamine-rich domain, CSD: cold shock domain, ncCSD: non-canonical cold shock domain, (Hollmann et al., 2020)). **C:** Schematic model of the full-length *msl-2* mRNA with reported binding sites for Sxl (green), Hrp48 (red) and Unr (blue). Sites B, E and F are required for optimal translational repression (Gebauer et al., 1999; Gebauer et al., 2003). The white circle marks the region we studied in addition to the earlier identified binding site. Region 5 is the sequence suggested previously as the msl-2 Hrp48 binding site (Szostak et al., 2018). It can self-associate through duplex formation in isolation (Figure S4C). Hrp48 *P-element* binding site shows high similarity to a region overlapping Region 5 and 6 of *msl-2* (grey). The 6-mer constructs were used in this study to refine the Hrp48 binding site. **D:** Isothermal titration calorimetry (ITC) data of Hrp48-RRM1 titrated with 6-mer-3 and 6-mer-4. **E:** A zoomed-in region of the ^15^N,^1^H-HSQC NMR titration experiments of ^15^N-labeled Hrp48-RRM1 and 6-mer-3 and 6-mer-4. **F:** Mapping of the chemical shift perturbation (CSP) data of the RRM1-6-mer-3 titration on the RRM1 crystal structure (left panel) and sequence (right panel). The threshold for mapping a CSP onto the structure was one standard deviation of the CSP values. **G:** MD-derived Hrp48-RRM1-RNA complex with 6-mer-3 RNA. The overall structure is surrounded by zoom-ins of single nucleotides being base-specifically recognized by RRM1 residues.

Despite advances in unraveling the complexity of this translational control mechanism, how Hrp48 interacts with *msl-2* 3’ UTR and represses translation is not well understood. The exact RNA recognition mode of Hrp48 and whether it establishes cooperative contacts with Sxl and Unr are not known.

In this study, we investigate RNA binding of Hrp48 in isolation and as part of a multiprotein complex involving Sxl and Unr bound to an extended site F *msl-2*-derived RNA fragment. We used NMR spectroscopy and molecular dynamics simulations to refine the cognate RNA motif within the 3’ UTR, which partially overlaps with the previously reported motif (Szostak et al., 2018). Furthermore, we determined binding affinities of the two structured RNA recognition motifs (RRM) of Hrp48, which jointly bind *msl-2*. NMR ^15^N spin relaxation and titration data suggest that the two domains bind RNA cooperatively and tumble together in the RNA bound state. *In vitro* translation assays support our findings on the binding site at the functional level. We report the crystal structure of Hrp48 RRM1 at 1.2 Å resolution and validate the structure prediction model of RRM2 by NMR spectroscopy. Further biochemical and biophysical data on Hrp48, Unr, Sxl and *msl-2* suggest no interaction of the proteins in the absence of RNA. Notably, the three proteins bind *msl-2* simultaneously and we established a protocol to reproducibly form the quaternary complex. Our results suggest that Hrp48 binds *msl-2* mRNA independently of the Sxl-Unr moiety in absence of its intrinsically disordered regions (IDRs).

## Results

### Structure and RNA binding of Hrp48-RRM1

Initially, we used the divide-and-conquer approach and investigated each RRM domain of Hrp48 in isolation with regard to their structure and RNA binding properties. To this end, we conducted isothermal titration calorimetry (ITC) and ^15^N,^1^H-HSQC NMR experiments to observe chemical shift perturbations of protein resonances upon titration with RNA. The previously reported *msl-*2 binding site of Hrp48 (AACCUAGG) is downstream and adjacent to the Unr binding site close to the Sxl F site and has been termed Region 5 (Figure 1C, (Szostak et al., 2018)). Overlapping with this Region 5 but shifted four nucleotides towards the 3’ end is a sequence (UAGGAUUAAG), which is highly similar to a sequence reported earlier to be the Hrp48 binding site on *P-element* mRNA (UAGGUUAAG, Figure 1C, (Siebel et al., 1992; Siebel et al., 1994). Thus, we divided the 17 nucleotides just downstream of the Sxl (F site)- Unr binding region into 6-mer RNA oligonucleotides to assess binding of the single RRM domains while preventing formation of double-stranded RNA (dsRNA) by the palindromic motif of Region 5 (Figure 1C). This resulted into four 6-mers with the following sequences: 6- mer-1: AACCUA, 6-mer-2: UAGGAU, 6-mer-3: AUUAAG and 6-mer-4: AAGAAC (Figure 1C). We also added a U_6_-mer as a control (6-mer-5, Figure S1), a sequence unrelated to the other 6-mers. In our ITC experiments, we tested RNA binding of RRM1 for each 6-mer. Binding of the control U_6_-mer (6-mer-5) was not strong enough to be detected by ITC. From ITC-derived dissociation constants (*K*_D_) the optimal motif for RRM1 could not be identified as they were in the same order of magnitude for all four 6-mers (Figure 1D, S1A-C, Table 3). Interestingly, a clear difference in the enthalpic and entropic contributions to the affinity of binding could be observed between 6-mer-1/4 and 6-mer-2/3 (Figure 1D and S1A, B Table 3). We then used NMR titration experiments to further study the RNA binding and complement the ITC results. Based on the CSPs, RRM1 binds to all five hexamers, including the control 6- mer-5, but with different affinities (Figure 1E and S1D–H). For the 6-mer-1, 6-mer-4 and 6- mer-5 titrations, CSPs indicate that RNA binding is in the fast exchange regime on the NMR time scale, whereas for 6-mer-2 and 6-mer-3 we could observe strong CSPs and intensity changes, characteristic for an intermediate-to-slow exchange regime (Williamson, 2013). CSPs in the intermediate-to-slow exchange regime do not allow for reliable determination of dissociation constants. However, slow exchange usually indicates stronger binding with low micromolar affinities (Kleckner and Foster, 2011). Thus, from the NMR data, one could qualitatively conclude that 6-mer-2 and 6-mer-3 are bound stronger by Hrp48 than 6-mer-1 and 6-mer-4, with 6-mer-3 featuring the highest number of peaks with CSPs in the slow exchange regime. This is exemplified by tryptophan W16 located in the loop region between β1 and α1 (Figure 1F) shows the highest CSP upon 6-mer-3 binding and is in the intermediate-to-slow exchange regime (Figure S1D-H). Aromatic residues located in this region are likely to be involved in RNA binding which already has been shown for the RRM domain of Fox-1 (Auweter et al., 2006a; Auweter et al., 2006b). Thus, we concluded from the NMR titrations that 6-mer-3 is the best Hrp48 binder since this ligand leads to a bound state that is closest to a single low energy complex structure due to the high number of CSPs in the slow-intermediate exchange regime. Also, 6-mer-3 induces the strongest CSPs over most residues (Figure S1I). Backbone chemical shift assignment of RRM1 enabled us to map the CSPs onto the crystal structure of RRM1 (Figure 1F) we determined at 1.2 Å resolution (PDB: 9EN7) using molecular replacement based on the hnRNP A1-RRM1 structure (PDB: 1HA1, (Shamoo et al., 1997)) since hnRNP A1 is a putative mammalian homolog of Hrp48 which has 62% sequence similarity and 43% sequence identity (Figure S2). The structure of Hrp48-RRM1 adopts the canonical βαββαβ topology with four β-strands folding into an antiparallel β-sheet which is packed against two α-helices (Figure 1F, (Nagai et al., 1990)). The CSPs show that the RNA binds along the four β-strands, resembling the canonical RNA binding mode of RRM domains. Interestingly, peaks corresponding to residues located C-terminal to β-strand 4 just before the linker (P81-R85) between RRM1 and RRM2 starts, exhibit also strong CSPs. These may be induced by direct RNA binding or by allosteric effects.

As we could not determine an experimental RRM1-RNA complex structure, we utilized unbiased atomistic simulations using a recently rescaled RNA force field that allows observing the full spontaneous binding process of single-strand RNAs to proteins (Krepl et al. 2022). We applied this method to study the binding of 6-mer-3 (UUAAG) to RRM1, generating a large structural ensemble (∼110 μs in total; see Methods). The calculated ensemble was subsequently visually analyzed and all the observed bindings annotated against the NMR CSP data. A single representative RRM1-6-mer-3 complex binding motif, which excellently reflected the CSP data while also demonstrating long-term stability in MD simulations, was finally selected for further analysis (Figure 1G). In this structure, direct protein-RNA contacts are established where U2 (numbering according to the appearance within the full length 6-mer-3) is sandwiched between the indole ring of W16 and guanidinium group of R75. The arginine is oriented in a way that also allows formation of base-specific hydrogen bonds or electrostatic interaction with the downstream phosphate group during thermal fluctuations. The U3 is also recognized base-specifically by D78 and K80, forming hydrogen bonds, whereas adenosine 4 stacks with F10 and is base-specifically recognized by hydrogen-bonding with the backbone carbonyl of P81. Adenosine 5 forms a base-specific hydrogen bond network with the backbone carbonyls of N83 and P84 while the base stacks with the guanidinium group of R85. G6 is also recognized base-specifically by hydrogen bonds with the side chain of D35 and K8 while V37 forms hydrophobic interactions with the purine ring. Thus, all bases of 6-mer-3 can be contacted in a base-specific fashion, providing the structural basis for Hrp48’s highly sequence-specific RNA binding.

### Biophysical characterization of Hrp48-RRM2

Next, we wanted to assess the structure and RNA binding properties of Hrp48-RRM2. Crystallization trials with or without RNA of RRM2 were not successful. Therefore, we used AlphaFold2 to predict the structure of RRM2 (Jumper et al., 2021). Interestingly, the predicted structure revealed an additional β-strand between the last α-helix and β-strand, deviating from the canonical βαββαβ fold of RRM2 (Figure 2A). Secondary structure prediction based on our NMR chemical shift assignment of RRM2 using TALOS+ (Shen et al., 2009) confirmed the presence of this additional β-strand (β4) in solution (Figure 2B). During purification and analysis of NMR data we observed that RRM2 tends to oligomerize. Size-exclusion chromatography coupled with multi-angle light scattering (SEC–MALS) confirmed that this isolated domain forms tetramers in an isolated *in vitro* context at 20 µM and above (Figure 2C). ITC data suffered from noise at these low concentrations, indicating generally lower binding affinity of RRM2 compared to RRM1. However, ITC could not be performed at higher concentrations due to the oligomerization tendency, thus we utilized NMR for titration experiments to assess the RNA binding specificity and affinity of RRM2. Upon titration with 6-mer-1 and control RNA 6-mer-5 no or only weak CSPs could be observed (Figure 2D and S3). Titration with 6-mer-3 and 6-mer-4 resulted in CSPs in the fast and fast-to-intermediate exchange regime, whereas 6-mer-2 titration induced CSPs in the intermediate-to-slow exchange regime (Figure 2D and S3). The latter could not be fitted to obtain a *K_D_* value but it qualitatively indicates that 6-mer-2 binds strongest to RRM2. Thus, for RRM2, 6-mer-2 is the best binder. Mapping the CSPs onto the RRM2 structural model shows that the canonical β-strands are involved, typical for RRM domains (Figure 2E). Interestingly, also peaks corresponding to residues of the non-canonical β4-strand and α2-helix exhibit strong CSPs upon titration with 6- mer-2. This might be due to direct interaction with RNA or due to allosteric effects.

**Figure 2:**
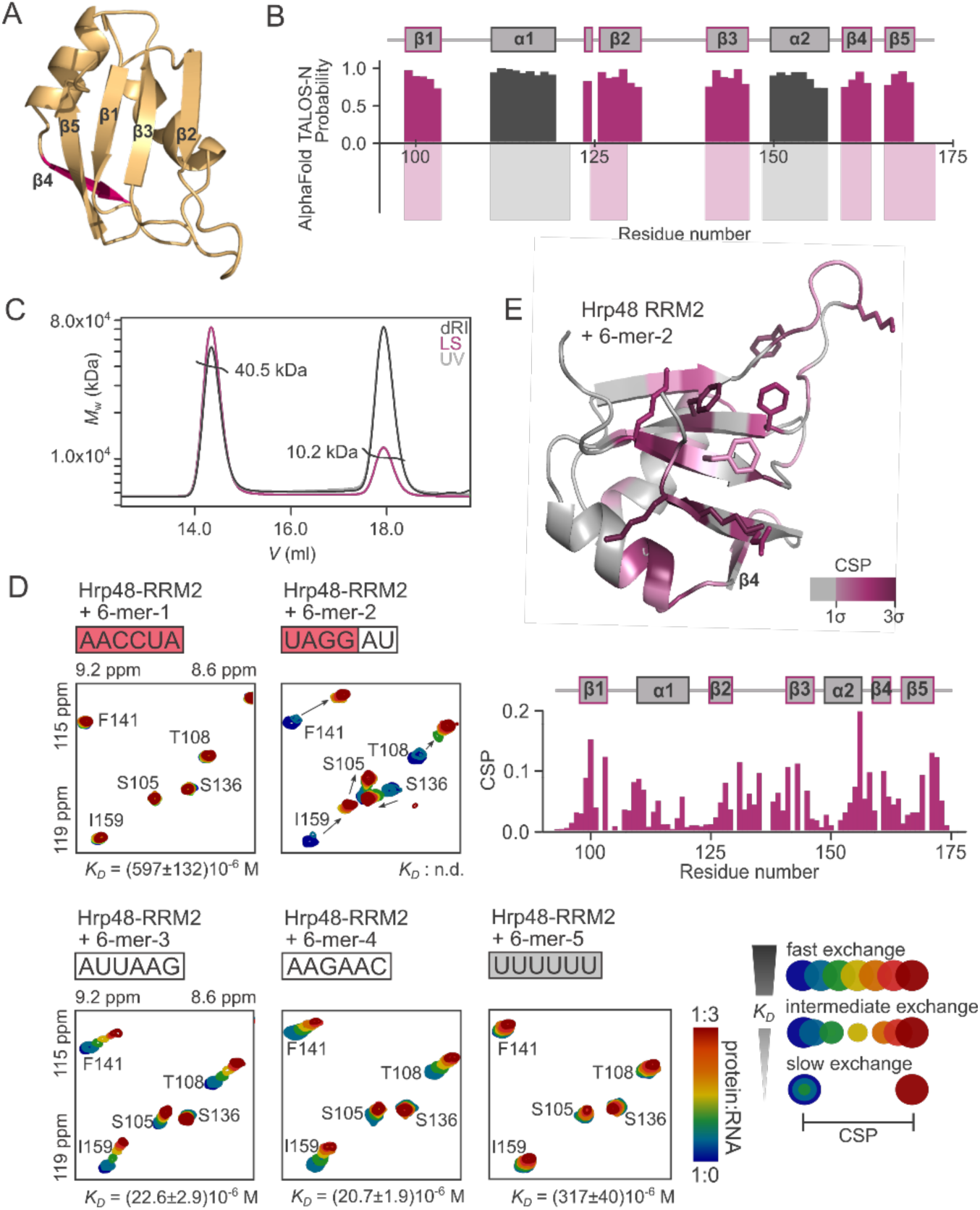
Structure and RNA binding of Hrp48-RRM2. **A:** AlphaFold2 (AF2) model of RRM2 features a non-canonical RRM fold with an additional β-strand, depicted in magenta (β4) (Jumper et al., 2021). **B:** The βαββαβ fold prediction of AF2 is confirmed by NMR chemical shift assignment data used in TALOS+ (Shen et al., 2009). **C:** Size exclusion chromatography coupled with multi-angle light scattering (SEC-MALS) of RRM2 reveals a strong oligomerization tendency of RRM2. Dark grey: differential refractive index (dRI), purple: light scattering (LS), light grey: UV absorption (UV). **D:** Zoomed-in regions of the ^15^N,^1^H-HSQC NMR spectra of ^15^N-labeled Hrp48-RRM2 titrated with the five different 6-mer RNA oligos to three-fold excess. RRM2 does not bind 6-mer-1 and weakly binds 6-mer-5. 6-mer-3 and 6-mer-4 binding is in the fast exchange regime on the NMR time scale, and 6-mer-2 in the intermediate-slow exchange regime. Schematic representation of CSP patterns indicating the different exchange regimes observable in NMR titrations. **E:** Mapping the CSPs on the structure of RRM2 reveals a mostly canonical RRM-RNA recognition mode.

We conclude that both RRM domains of Hrp48 recognize *msl-2* and prefer binding to 6-mer-2 and 6-mer-3 which together embrace a 10-mer recognition sequence (Figure 3A). RRM2 binds upstream of RRM1 which is also consistent with the binding behavior of RRM tandem domains of other proteins (Deo et al., 1999; Handa et al., 1999; Schäfer et al., 2019). Furthermore, the recognition site of Hrp48 on the *msl-2* mRNA is very similar to the Hrp48 binding site on *P-element* mRNA which strengthens the validity of our results (Siebel et al., 1992; Siebel et al., 1994). Next, we assessed binding of the tandem RRM12 construct of Hrp48 to *msl-2* mRNA.

**Figure 3:**
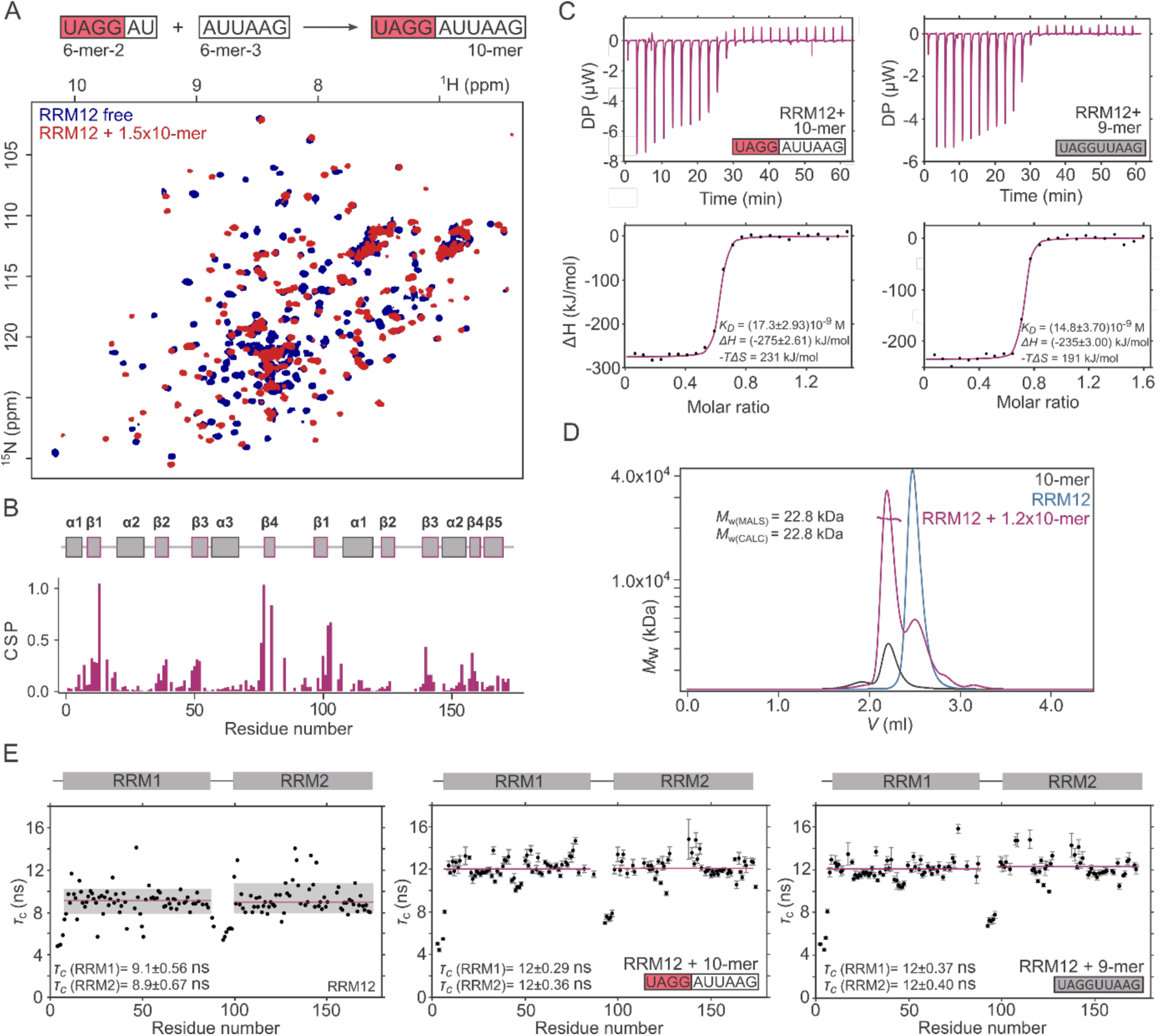
RNA binding and dynamics of Hrp48-RRM12. **A:** 10-mer RNA consisting of 6-mer-2 and 6-mer-3 sequences (upper panel) was used in ^15^N,^1^H-HSQC NMR titration experiment for RRM12 (lower panel). Peaks are in the slow-exchange regime; additional titration points are shown in Figure S4A. **B:** CSP plot of RRM12-10-mer NMR titration. Both RRM domains are involved in the interaction and residues in the β-strands are the most affected, including β4 of RRM2. **C:** ITC of Hrp48-RRM12 with 10-mer (left panel) and 9-mer (right panel), respectively. **D:** The 10-mer and RRM12 forms a 1:1 complex. SEC-MALS UV absorption chromatograms of free RRM12 (grey), 10-mer (blue) and their complex (purple). **E:** Rotational correlation times of RRM12 residues in the apo state (left panel), bound to 10-mer (middle panel) and bound to 9-mer (right panel). The domains tumble jointly in an RNA bound state. Left panel: grey: one standard deviation of the mean value of τ_c_ of each residue within a single RRM domain. Purple: Mean value of τ_c_ of residues within the standard deviation. Values outside the standard deviation are considered as outliers and were excluded for calculating the mean.

### The tandem domain RRM12 of Hrp48 binds *msl-2* cooperatively

For binding studies of the tandem domain RRM12 of Hrp48, we used 10-mer RNA (UAGGAUUAAG, Figure 3A). This sequence is four nucleotides downstream of the start of the previously reported Region 5 (Szostak et al., 2018), and consequently does not include the palindromic motif, allowing meaningful *in vitro* investigations without the risk of forming dsRNA. Indeed, we could confirm self-complementary base pairing of Region 5 by 1D-NMR which showed strong peaks in the imino region, indicative of base pairing (Figure S4A). With this 10-mer RNA, we wanted to test whether a tandem RRM12 construct would bind stronger and performed an NMR titration and ITC experiments to determine the binding affinity (Figure 3A-C). In the NMR titration, resonances corresponding to residues of both domains exhibited strong CSPs in the slow exchange regime indicating strong binding in the nanomolar range (Figure 3A, S4B). Both domains exhibit considerably larger CSPs upon RNA binding than the isolated domains. This is confirmed by ITC experiments from which we could determine a dissociation constant of 17.3 ± 2.9 nM. Thus, RRM12 binds around 100-times stronger to this 10-mer RNA than the individual domains to their best 6-mer motifs. As such, both RRM domains bind to single-stranded RNA simultaneously and synergistically. We could also show that RRM12 forms a 1:1 complex with the 10-mer in solution by SEC-MALS (Figure 3D, S4C), which together with the NMR data confirms that both domains bind one RNA 10-mer.

Having identified the optimal RNA binder for RRM12, we attempted to obtain a crystal structure of an Hrp48-RRM12-RNA complex. However, as for the isolated RRM domains, we could not obtain crystals with RNA despite extensive crystallization trials. Instead, we assessed the dynamics of the tandem domain through NMR ^15^N spin relaxation experiments. In the absence of RNA both domains have similar global rotational correlation times τ_c_ of 9.1 ± 0.6 ns for RRM1 and 8.9 ± 0.7 ns for RRM2. This indicates that both domains tumble mostly independently from each other (Figure 3E), as it can be estimated that the rotational correlation time is approximately 0.6 times the molecular weight of a globular domain (Rossi et al., 2010). In case of fully independent tumbling of both domains, a τ_c_ of around 6 ns would be expected. Thus, the short linker influences the tumbling of each domain or weak unspecific interactions between both domains could lead to an elevated rotational correlation time. The linker residues exhibit much lower rotational correlation times, indicating a high degree of flexibility, which does not change upon addition of 10-mer RNA. Thus, the linker is not involved in RNA binding. However, in presence of RNA the rotational correlation times of both domains are 12 ± 0.3 ns for RRM1 and 12 ± 0.4 ns for RRM2. This clearly shows that both domains tumble together and do not move freely with respect to each other but adopt a fixed orientation in RNA-bound state. Consistent with the NMR titration data of the isolated RRM1 domain, we observed elevated τ_c_ values and significant CSPs for assigned residues located in between β-strand 4 and the linker residues, confirming that these residues are not flexible but involved in RNA contacts.

Since the Hrp48 binding site of *msl-2* mRNA is not identical to the recognition sequence on the *P-element* mRNA, we investigated whether the difference (one additional A in the center part of the *msl-2* mRNA binding sequence, UAGG**<u>A</u>**UUAAG) has an impact on the dynamics of the tandem domain RRM12 in the RNA bound state. Given that NMR relaxation data for both complexes resulted in almost identical rotational correlation times per residue, we conclude that the additional adenosine of the 10-mer does not lead to increased flexibility between both RRM domains (Figure 3E).

To test whether the additional adenosine has any impact on the binding affinity of the RRM domains towards the RNA we conducted ITC experiments by titrating 9-mer RNA to RRM12 (Figure 3C, right panel) and in addition used a 10-mer mutant sequence in which the two nucleotides in the center part of the 10-mer wild-type (10-mer-WT) are mutated to CC (UAGG**<u>CC</u>**UAAG, 10-mer-CC, Figure S4D). The *K_D_* value resulting from the 9-mer titration is with 14.8 ± 3.7 nM in the same range as for the 10-mer-WT experiment (17.3 ± 2.9 nM) which is consistent with the NMR relaxation data where the additional A does not induce changes in backbone dynamics of Hrp48. Interestingly, the ITC experiment for the 10-mer-CC titration suffered from severe noise, hence calculation of the binding affinity was not feasible (Figure S4D, right panel). However, since the experimental setup was identical to 10-mer-WT and 9-mer titration experiments, we conclude that the RRM12 has a significantly weaker affinity for 10-mer-CC than for the 10-mer-WT. However, it must be considered, that the 10- mer-CC is partly self-complementary and could form dsRNA, which would interfere with Hrp48 binding. Nevertheless, the result is consistent with our MD-derived structural model in which the central U (Figure 1G, U2 in 6-mer-3) is specifically recognized by R75. Substitution to a C would abolish these specific contacts and weaken the affinity.

Accordingly, we hypothesized that the adjacent U (U6 in 10-mer-WT) downstream of the additional A (A5 in 10-mer-WT) could have an important role for Hrp48-RRM12 binding. To validate this hypothesis, we used shortened RNA constructs of a length of 4 nucleotides (4-mer- 1 – 4-mer-6, Figure S5A) for NMR titration experiments. All 4-mers together encompass the entire 10-mer sequence and the previously identified Region 5 (Figure S5A). The results confirmed that only 4-mer-3 and 4-mer-6, which correspond to the 5’ and the 3’ end of our already confirmed 10-mer-WT binding sequence show stronger CSPs in comparison to all other 4-mer NMR titration experiments (Figure S5B, C). 4-mer-4 and 4-mer-5 that consist of sequences which include the central A and U, show lower CSPs compared to 4-mer-3 and 4- mer-6 which indicates that the central region binds weaker. We then used 4-mer-3 and 4-mer- 6 for NMR titrations to the single RRM domains to check whether a 4-nucleotide sequence is sufficient for efficient RNA binding (Figure S6A-E). CSP analyses revealed that RNA still binds both RRM domains but the affinity decreased significantly compared to 6-mer binding (Figure S6F). From these data we conclude that the RRM domains mainly bind to the 5’ and the 3’ end of the 10-mer-WT binding site, whereby four nucleotides are not sufficient for efficient binding, thus the central U is bound by Hrp48 whereas the central A is dispensable.

### Validation of Hrp48’s RNA binding by mutational analysis in translation assays

To further validate the refined Hrp48 binding site of *msl-2* mRNA, we performed a mutational analysis and tested the ability of the mutant mRNAs to be repressed by Sxl using translation assays in *D. melanogaster* embryo extracts. Several RNA constructs were tested for *in vitro* translation activity in Luciferase reporter assays (Figure 4, methods). We used CU-repeats to mutate specific parts of the 3’ UTR including 5m, 6m or fragments within (Figure 4A). We also investigated the impact on translation repression upon substituting the entire region (Region 5 and 6, designated 56m). The latter has the strongest de-repression effect compared at a RBD4/RNA concentration ratio of 10 (Figure 4A). Mutation of the 10-mer sequence (10m) derepressed translation to a lesser extent compared to 56m but still had an important effect compared to the wild-type sequence. Interestingly, mutation of the last 6 nucleotides of the 10- mer sequence (6m) resulted in similar translation de-repression than for the whole 10-mer, indicating that Hrp48-RRM1 is more important for translation repression than binding of RRM2. This hypothesis is supported by the results of the 10.1m luciferase assay. Here, translation repression of the reporter is as efficient as in the wild-type sequence. In addition, CSPs of the NMR titration experiments of RRM1 and RRM2 with 6-mer RNA indicate generally stronger binding of RRM1 compared to RRM2 which would also strengthen the aforementioned hypothesis. The translation repression assay with 10.3m resulted in a slightly de-repressed translation. However, this weak change in translation efficiency compared to the wild-type indicates that the central part of the 10-mer is not essential for translation repression. Mutating the complete sequence (56m), in which both adenosines at the 5’ end were also exchanged with a CU-repeat, had the largest translation de-repression effect. These two adenosines are in contact with Sxl which would explain the elevated translation de-repression effect for 5m and 56m (Hennig et al., 2014). Thus, we confirmed that the 10-mer motif is the optimal Hrp48 binding site and its mutation has an impact on translation repression.

**Figure 4:**
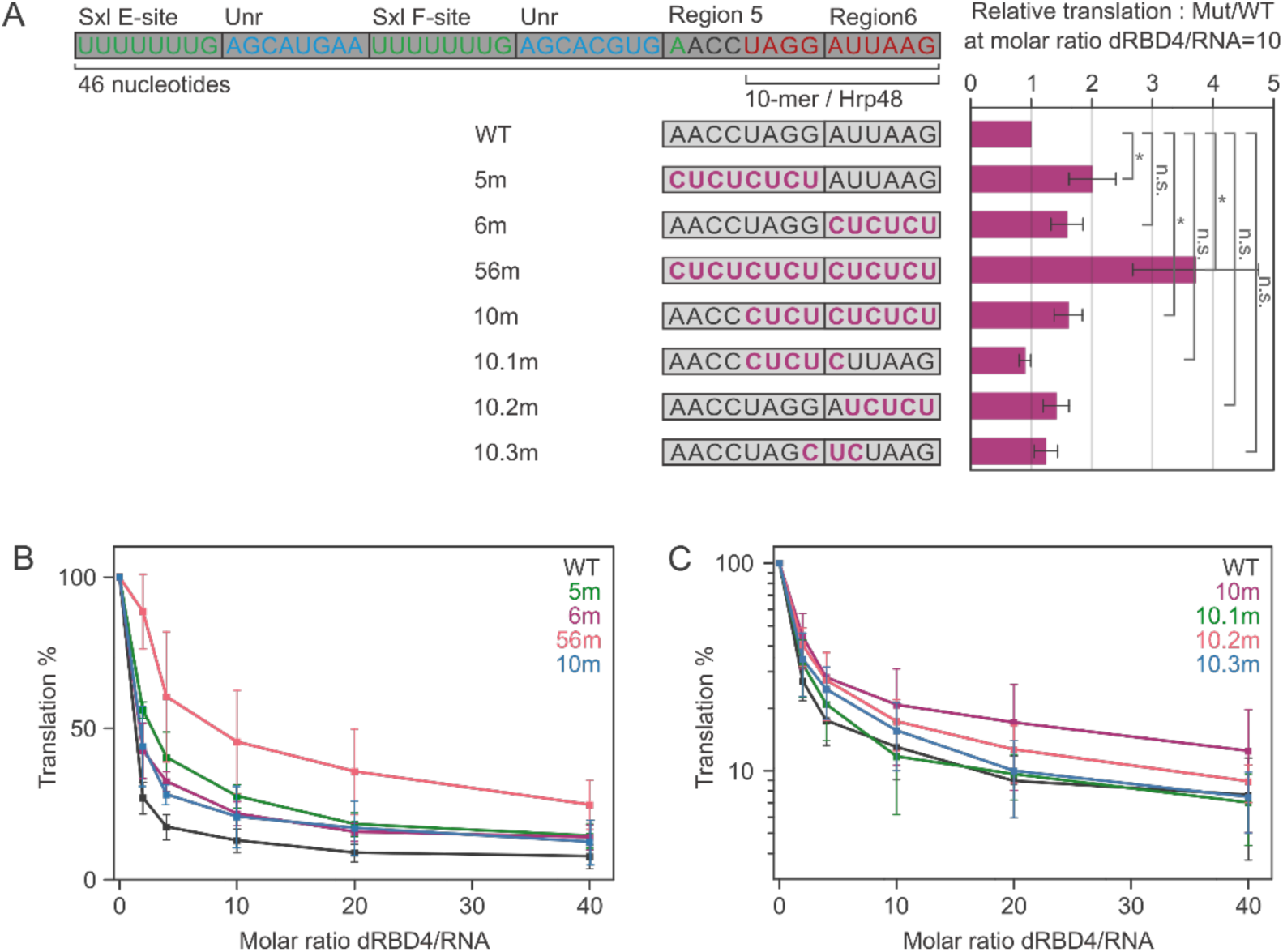
The Hrp48 RNA-binding site is necessary for translation repression. **A:** *In vitro* translation assays were performed with Firefly luciferase reporters appended to a segment of the 3’ UTR of *msl-*2 containing the minimal Sxl (green) and Unr (blue) binding sites, and derivatives of the Hrp48-binding region. The CU-repeat substitutions are indicated with bold pink fonts. The bar graph shows the relative translation of the luciferase reporter at a Sxl/RNA concentration ratio of 10. Co-translated Renilla luciferase was used as an internal control and as a reference for normalization of the reporter Firefly luciferase signal, and the data was referred to the wild type (WT) signal activity. Data represent the average of three independent experiments; error bars represent standard deviation. *p < 0.05. **B–C:** *In vitro* translation assay results for all Sxl/RNA concentration ratios used in the experiment. The data was plotted as relative translation taking as 100% the initial activity with no Sxl added to the extract.

### Complex formation of Hrp48 with Sxl and Unr is RNA dependent

Having obtained mechanistic insight into the Hrp48 binding mode and base specificity, we wanted to assess whether Hrp48-RRM12 forms a complex with Sxl and Unr and whether Hrp48-RRM12 binds to an extended RNA, including the F site of Sxl and Unr binding sites. However, we tested first whether Hrp48 interacts with Sxl or Unr in the absence of RNA. It has been shown previously that Sxl-dRBD3 and Unr-CSD1 do not interact without RNA, but protein-protein contacts between the two proteins are established upon RNA binding (Hennig et al., 2014). We expressed and purified unlabeled Sxl-dRBD3 and different constructs of Unr. The domain boundaries of Unr were based on previous work in our lab, where four additional non-canonical CSDs were identified in between the five predicted canonical CSDs (Figure 1B, (Hollmann et al., 2020)). ^15^N,^1^H-HSQC spectra were recorded of Hrp48-RRM12 alone, and Hrp48-RRM12 together with one equivalent Sxl or Unr (Figure S7A-D). We also performed the reverse experiment with labeled Sxl-dRBD3 and unlabeled Hrp48-RRM12 (Figure S7E). Due to the lack of chemical shift perturbations or significant intensity decrease in all titrations, we concluded that Sxl-dRBD3 and the tested Unr constructs do not interact with Hrp48-RRM12 in the absence of RNA. The small decrease in intensity for some of the peaks might be a result of unspecific interactions or aggregation of the proteins. Overlay of SEC–MALS chromatograms of the separate proteins and all three proteins together further confirmed that the constructs we later used for complex formation, Sxl-dRBD3, Unr-CSD12 and Hrp48- RRM12 do not interact in the absence of RNA (Figure 5A).

**Figure 5:**
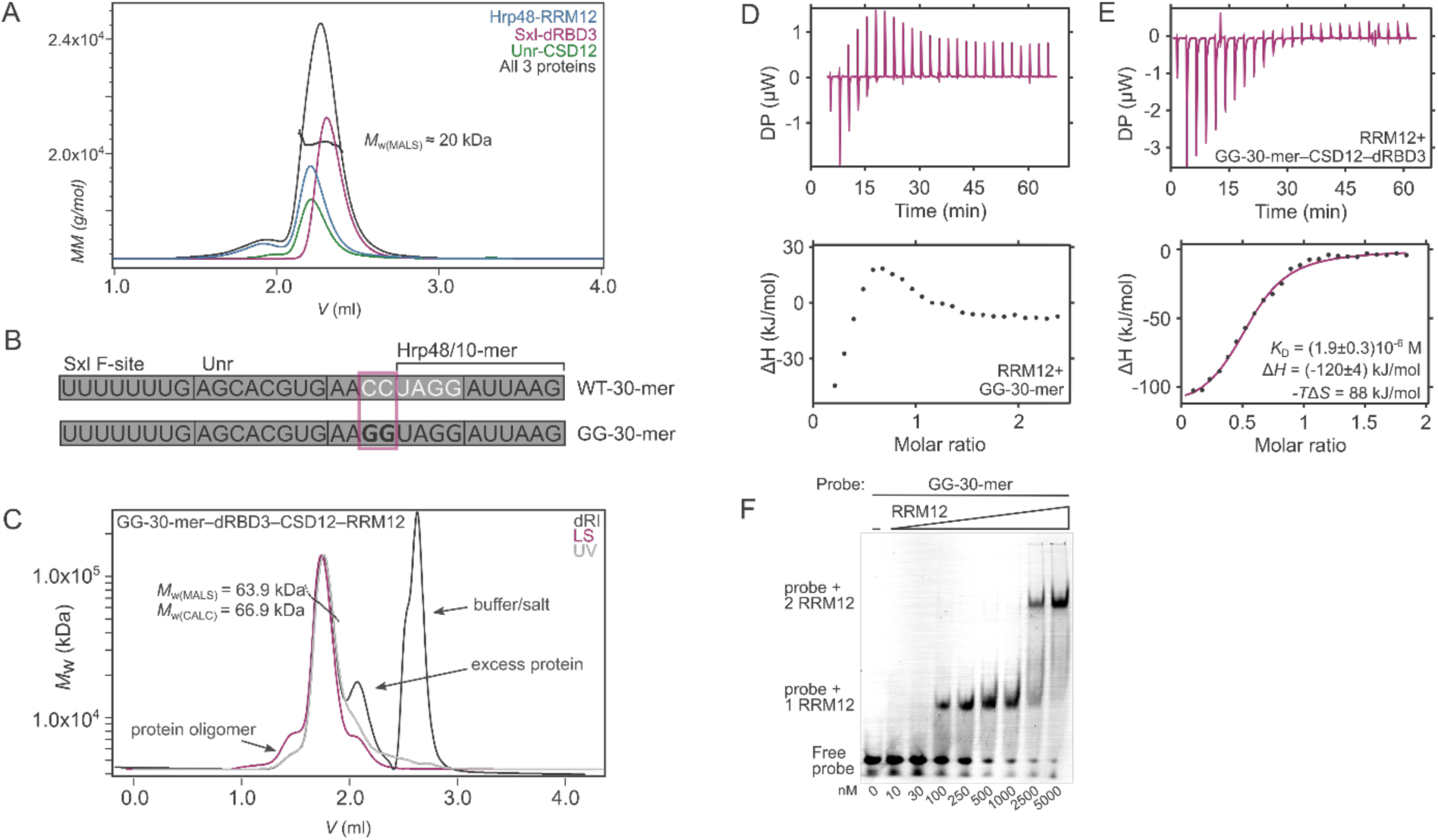
Complex formation of Hrp48, Sxl, Unr and RNA. **A:** Hrp48-RRM12, Sxl-dRBD3 and Unr-CSD12 do not interact in the absence of RNA. SEC–MALS UV absorption chromatograms of the proteins Hrp48-RRM12 (blue), Sxl-dRBD3 (purple), Unr-CSD12 (green) and the proteins at the same concentration injected at once (grey). The molecular weight is around 20 kDa, which corresponds to the *M*_w_ of the free protein constructs. **B:** The RNA constructs used for complex formation. Both constructs contain one binding site for each protein. The GG-30-mer is a mutant in which the palindromic sequence is abolished by substituting the indicated (framed) CC into a GG pair to decouple the dimerization from protein binding in biophysical experiments. We also confirmed that this substitution has no effect on translation repression (Figure S7F). **C:** SEC-MALS of the GG-30-mer-Hrp48-RRM12-Unr-CSD12-Sxl-dRBD3 complex. (LS), dark grey: differential refractive index (dRI), purple: light scattering, light grey: UV absorption. The molecular weight was calculated for a 1:1:1:1 complex. **D – E:** ITC of GG-30-mer titrated into Hrp48-RRM12 (D) and of GG-30-mer-Unr-CSD12-Sxl-dRBD3 titrated into Hrp48-RRM12 (E). **F:** Electrophoretic mobility shift assay using the GG-30-mer and increasing amounts of Hrp48-RRM12. At higher concentration of Hrp48 an additional shift appears, indicating that two Hrp48-RRM12 molecules bind one RNA.

With the intention of testing whether Sxl, Unr and Hrp48 jointly interact with *msl-2*, which has been proposed earlier (Szostak et al., 2018), we designed two 30-mer RNA constructs (WT-30- mer and GG-30-mer) based on published data for Sxl and Unr and our findings of the Hrp48 binding site (Hennig et al., 2014) (Figure 5B). This site combines the F site to which Sxl binds, followed by the Unr and the 10-mer Hrp48 binding site. In the 30-mer-GG, we introduced a mutation to avoid the palindromic sequence and preclude formation of dsRNA (Figure 5B). A negative effect of this mutant on the functionality was not detectable in translation assays (Figure S7F). For complex formation studies involving RNA, we chose shortened constructs of Sxl (Sxl-dRBD3), Unr (Unr-CSD12) and Hrp48 (Hrp48-RRM12), that we know interact with *msl-2* mRNA.

To determine the best conditions for complex formation, we injected the two proteins Sxl- dRBD3 and Unr-CSD12 together with the GG-30-mer, with different molar ratios. We obtained the highest amounts of complex at a 1:2:3 ratio (RNA:dRBD3:CSD12, Figure S7G). Then we optimized the Hrp48 amount, and accordingly we used two-fold excess with respect to the RNA (Figure 5C). The resulting main peak of the chromatogram corresponds to the complex. The discrepancy of the calculated and determined molecular weight could be explained by low resolution and similar retention times of the overlapping peaks (Figure 5C). While pre-forming the ternary complex with WT-30-mer in absence of Hrp48-RRM12, we found that the molecular weight is almost double of what was expected (Figure S7I). This could be explained by dsRNA formation due to the palindromic sequence located within the WT-30-mer. For this reason, we used GG-30-mer in the ITC and electrophoretic mobility shift assays, to avoid additional interactions complicating the biophysical characterization of the complex. However, upon addition of Hrp48-RRM12, we could detect the monomeric complex, thus binding of Hrp48-RRM12 resolves the formed dsRNA structure of the WT-30-mer and forms a stable quaternary complex with 30-mer RNA, Sxl-dRBD3 and Unr-CSD12 (Figure S7H, I).

Next, we wanted to assess whether Hrp48-RRM12 binding to RNA is synergistic with Sxl- dRBD3 and Unr-CSD12 binding to 30-mer RNA. To this end, we performed ITC experiments. GG-30-mer titrated into Hrp48-RRM12 provided a complex, but reproducible ITC curve, which could indicate binding of multiple RRM12 domains on the long GG-30-mer (Figure 5D). Another putative binding site of RRM12 on the GG-30-mer could be the Sxl F binding site, since our data of the 6-mer NMR titrations for RRM1 showed also weak interactions to the U_6_ RNA (6-mer-5, Figure S1E). However, it cannot be excluded that RRM12 could also bind to another part of the RNA. To check for multiple binding sites for Hrp48 on the GG-30-mer, we performed an electrophoretic mobility shift assay (EMSA), which resulted indeed in two sequential binding events (Figure 5F). The first shift indicates binding of one Hrp48-RRM12 domain to the GG-30-mer and the second shift suggests that multiple Hrp48-RRM12 proteins could be bound to the RNA. We consider the second binding event as biologically irrelevant and as an *in vitro* artefact, as the affinity is very weak and this site is mostly occupied by Sxl and Unr. However, this could explain the complex ITC curve we obtained (Figure 5D). ITC titrations of a pre-formed complex of GG-30-mer, Sxl-dRBD3 and Unr-CSD12 titrated into Hrp48-RRM12 abolished the second binding event and we could measure the *K_D_* of Hrp48- RRM12 binding to this preformed complex. To our surprise, the affinity was about 100-fold weaker than when Hrp48-RRM12 interacts with the 10-mer RNA, which at the very least rules out synergistic binding of Hrp48-RRM12 to GG-30-mer in the presence of Sxl-dRBD3 and Unr-CSD12. However, it cannot be excluded that Hrp48’s C-terminal IDR interacts with Sxl, Unr or both proteins. We could not include this IDR in our current studies as it mediated severe aggregation.

## Discussion

In summary, we identified the binding site for the individual RRM domains of Hrp48 by mapping the region of *msl-2* using 4-mers and 6-mers. Since the binding affinity upon 4-mer interaction decreased significantly compared to 6-mer binding, we concluded that the smallest possible binding motif for the single domains must be at least five nucleotides long. Furthermore, the recognition sequence for each RRM domain of Hrp48 includes a motif highly similar to the consensus binding sequence (UAGG) of RRM1 of hnRNP A1, a putative mammalian homolog to Hrp48 (Beusch et al., 2017; Burd and Dreyfuss, 1994). The crystal structure also shows the characteristic βαββαβ fold for Hrp48-RRM1, whereas RRM2 possesses an additional β-strand, which is also involved in RNA binding. This non-canonical RRM fold has also been observed for PTB RRM3 where it extends the length of the β-sheet to accommodate more nucleotides and thus increase sequence-specificity (Conte et al., 2000). This could be similar for Hrp48, as CSPs were observed upon RNA binding for residues of this additional β-strand. Furthermore, the binding affinity of RRM2 towards 4-mer RNA decreases more than for RRM1, which could also indicate that more than four nucleotides are bound on the extended β-sheet of RRM2. On the other hand, CSPs on the additional β-strand could also occur due to allosteric effects. Additional β-strands have been reported for both RRM domains of TDP-43 which promote protein–protein interactions (Kuo et al., 2009). Other secondary structure elements as addition to the canonical RRM fold have been shown to increase plasticity and RNA specificity of RRMs (e.g. (Cléry et al., 2008; Duszczyk et al., 2022; Wang et al., 2014)). Preliminary data suggests that the self-interaction of Hrp48 through its RRM2 domain diminishes upon RNA binding, and that the RNA binding interface of RRM2 overlaps with its tetramerization interface. This might suggest a biological relevance of tetramerization, which might prevent unspecific RNA binding as in a tetramer the RNA binding interface of RRM2 is potentially buried. Dimerization or oligomerization tendencies of RRM domains to be biologically relevant has been shown for several other examples (e.g. (Mackereth et al., 2011; Pabis et al., 2019; Ripin et al., 2019)) The tetramerization tendency might also be a step in formation of higher oligomeric assemblies and eventually phase separation, as it has been shown in the context of *P-element* mRNA localization (Bose et al., 2022).

Both RRM domains of Hrp48 bind to 6-mer RNA motifs adjacent to each other, hence suggesting a continuous binding site of 10 nucleotides for the tandem RRM12 domain. The high affinity, simultaneous binding of RRM12 and the high similarity to the *P-element* mRNA binding site (Siebel et al., 1992; Siebel et al., 1994) also support the hypothesis of 10-mer RNA (UAGGAUUAAG) being the optimal sequence for Hrp48 binding on *msl-2*-mRNA. Additionally, RRM2 binds upstream of RRM1 which is also consistent with the binding behavior of RRM tandem domains of other proteins (Deo et al., 1999; Handa et al., 1999; Schäfer et al., 2019). Studies about dynamics revealed that both RRM domains which are connected through a short seven nucleotide linker do not interact with each other in absence of RNA but tumble jointly in an RNA bound state. Experimental data strengthen this hypothesis, since 9-mer RNA, representing the *P-element* mRNA binding site which does not contain the central adenosine, has equal binding affinity to RRM12 as the 10-mer. Furthermore, the adenosine does not provide additional flexibility, since rotational correlation times of RRM12 bound to 10-mer-WT and 9-mer RNA are identical.

Activity assays confirmed that upon mutation of the binding site, translation is partially derepressed which supports the validity of the binding site in a functional aspect and that Hrp48 contributes to translation repression of *msl-2*. In context of complex formation, Hrp48-RRM12 does not interact with Sxl and Unr in absence of RNA and also does not establish cooperative protein-protein contacts when bound to RNA. However, the results of complex formation studies only provide information about the RNA binding domains of the three proteins, thus, potential protein-protein interactions are still possible if they are established through contacts between Hrp48 IDRs or with other CSDs of Unr. A similar situation has been observed for the quaternary protein-RNA complex of Brain tumour (Brat), Pumilio (Pum) and Nanos, which together repress the translation of *hunchback* mRNA. This is an important step during *Drosophila* development, regulating the establishment of the anterior-posterior body axis. While the Pum-HD-Nanos-ZnF and Brat-NHL domains bind to adjacent RNA sequences (Loedige et al., 2014; Loedige et al., 2015; Weidmann et al., 2016), only Pum-HD and Nanos-ZnF exhibit cooperativity, whereas Brat-NHL binds independent of the Pum-HD-Nanos-ZnF module to RNA (Macošek et al., 2021). However, also in this case it cannot be excluded that the IDRs of each protein would engage in further cooperative contacts. Despite the experimental difficulties often associated with IDRs (aggregation and precipitation), it will be essential for future studies to include these regions to obtain a complete picture of these complexes’ mechanisms of action.

For the complex studied here, it needs to be considered that other proteins are also involved and could engage in further protein-protein contacts. Especially poly(A)-binding protein (pAbp), which promotes closed-loop formation of the mRNA (Vicens et al., 2018) and translation has been shown to take part in the *msl-2-*mRNA translation repression mechanism by interacting with the Sxl-Unr complex (Duncan et al., 2009). The interactions between Unr and pAbp are direct (Hollmann et al., 2023) but the molecular mechanism behind pAbp involvement remains elusive.

The Sxl-Unr-Hrp48 and the Pum-Nanos-Brat complexes are promising model systems to obtain insights into regulation of translation initiation and we are optimistic that cryo-EM or integrative structural biology (Dimitrova-Paternoga et al., 2020) will enable high-resolution structure determination of such full-length complexes in the future.

## Materials and Methods

### Protein cloning, expression and purification

The sequences of Hrp48 (UniProt code P48809), RRM1 (1 – 88), RRM2 (89 – 173), and the tandem RRM12 (1 – 173) constructs were cloned into the pETM-vector using restriction-free cloning (van den Ent and Löwe, 2006). Hrp48 constructs were expressed in *Escherichia coli* BL21 (DE3). Expression and purification of Sxl-dRBD3 (123-294) and all Unr constructs used in this study are described elsewhere (Hennig et al., 2014; Hollmann et al., 2020). Proteins were expressed in TB and LB medium or in M9 minimal medium for isotope labeling using ^15^NH_4_Cl for ^15^N labeled samples, or ^15^NH_4_Cl and D-Glucose-^13^C_6_ for ^15^N, ^13^C labeled samples. Generally, precultures were grown in the same medium as the expression medium overnight at 37°C. Cultures were inoculated with an OD_600_ of 0.04 and grown until the logarithmic growth phase was reached, followed by induction with 0.2 mM IPTG and overnight expression at 18°C. The harvested cells were resuspended in ice cold lysis buffer (20 mM Hepes, 500 mM NaCl, 12 mM imidazole, 0.5 mM β-mercaptoethanol, pH 7.2) and then frozen and sonicated for further lysis. The thawed lysate was centrifuged for 30 mins at 18000×g at 4°C and the supernatant was syringe-filtered with 0.45μm pore size filter. The cleared lysate was loaded three times on a Nickel-nitrilotriacetic acid (Ni-NTA) gravity flow column that was pre-conditioned in binding buffer (20 mM Hepes, 500 mM NaCl, 12 mM imidazole, pH 7.2). followed by adding the cleared lysate, washing with 20 column volumes of binding buffer and elution with 5 column volumes of elution buffer (20 mM Hepes, 500 mM NaCl, 200 mM Imidazole, pH 7.2). The His_6_-tag was cleaved by TEV protease and 2 mM β-mercaptoethanol, incubated on ice for 1 hour, and then dialyzed overnight at 4°C into binding buffer to remove the β-mercaptoethanol. The sample was reloaded onto the Ni-NTA gravity flow column collecting the flow-through to remove the cleaved tag. For Unr CSD1, CSD12, and CSD123, the reverse Ni-affinity step was followed by a heparin purification. For this, the flow-through was diluted five times to lower the concentration, and then dialyzed against the low salt heparin binding buffer (10 mM MES, 50 mM NaCl, 2 mM DTT, pH 6.0) overnight. The protein was injected onto a HiTrap^TM^ 5 mL Heparin HP, washed with heparin binding buffer and eluted with a heparin elution buffer (20 mM MES, 1500 mM NaCl, 2 mM DTT, pH 6.0) to remove bacterial RNA contamination. All proteins were further purified by size-exclusion chromatography (SEC) on a HiLoad 16/600 Superdex S75 or S200 pg column equilibrated with SEC buffer (20 mM MES, 200 mM NaCl, pH 6.5). Hrp48 RRM2 and RRM12 were kept at low concentrations (0.1-0.2 mg/ml) during all steps after the second Ni-NTA purification step to avoid aggregation.

For Sxl-Unr-Hrp48-*msl-2* complex formation, the purified Sxl-dRBD3, Unr-CSD12 or CSD1 and Hrp48-RRM12 constructs and the WT-30-mer or GG-30-mer RNA were incubated on ice in a 2:3:2:1 ratio. The complex was concentrated with a 3.5 kDa cut-off concentrator unit. Depending on the required quantity, the final volume was ∼300 μL for SEC purification on a Superdex 75 10/300 GL column, or ∼1 ml for the HiLoad 16/600 Superdex 75 pg column. The identity of the complex peak was confirmed by UV absorption measurement at 260 and 280 nm and also using SEC-MALS weight determination.

After SEC, the pooled fractions were concentrated and the concentration was measured at 260 nm on a NanoDrop UV-VIS absorption spectrophotometer with an extinction coefficient of 325000 (M^-1^cm^-1^), that is the sum of the extinction coefficients of the components at 260 nm.

### Isothermal titration calorimetry

ITC measurements were performed using a Malvern Panalytical MicroCal PEAQ-ITC calorimeter at 20°C. For all experiments, the protein was loaded into the cell and RNA or protein–RNA complex into the syringe (Table 3). Prior, the samples were dialyzed against MES buffer (20 mM MES, 200 mM NaCl, 0.02% NaN_3_, pH 6.5) buffer overnight, adjusted to the appropriate concentrations, centrifuged at 15000 rpm for 10 min, transferred to the final tubes for measurements and degassed for 5 min. The titrations were accomplished with 19 – 25 injections corresponding to 1.5–2 μl injection volumes. Each injection lasted 3 seconds followed by a 150 second delay. The sample stirring was set to 750 rpm with reference power was 10 μcal/s. Further details about the concentrations and experimental setup in individual titrations are listed in Table 3. The software MicroCal PEAQ-ITC analysis software (Malvern Panalytical) was used for data analysis.

### NMR spectroscopy

All NMR spectra were recorded on Bruker Avance III NMR spectrometers with magnetic field strengths corresponding to proton Larmor frequencies of 600 MHz, 700 MHz, 800 MHz or 1 GHz equipped with a room temperature triple resonance probe head (700 MHz), or a cryogenic triple resonance gradient probe head (600, 800 MHz and 1 GHz) at 298 K. The NMR samples were measured in SEC buffer with 5% D_2_O for the deuterium lock. The multidimensional experiments were recorded using apodization weighted sampling (Simon and Köstler, 2019). Backbone resonance assignment of ^13^C, ^15^N-labeled Hrp48 constructs was achieved to a completion of 100% (RRM1, excluding prolines), 94% (RRM2), and 95 % (RRM12) in the free state and 79% in the 10-mer bound state using ^1^H,^15^N-HSQC, HNCO, HN(CA)CO, HNCA, CBCA(CO)NH, HNCACB triple resonance experiments (Sattler et al., 1999). All NMR spectra were processed using NMRPipe (Delaglio et al., 1995), analysed using CcpNmr Analysis (Skinner et al., 2016; Vranken et al., 2005), CARA (Keller, 2004), Sparky (Lee et al., 2015) and NMRViewJ (Johnson, 2004). Backbone torsion angles were predicted from C_α_ and C_β_ chemical shifts for RRM2 using TALOS-N (Shen et al., 2009). The NMR backbone chemical shift assignments are deposited at the Biological Magnetic Resonance Databank under the following accession codes: 52354 (Hrp48 RRM2), 52359 (Hrp48 RRM1), 52362 (Hrp48 RRM12 bound to 10-mer-WT RNA).

NMR titrations were performed by recording two-dimensional ^15^N,^1^H-HSQC spectra of the labeled protein and for each subsequent addition of the titrant protein or RNA, until reaching saturation (i.e. no further chemical shift perturbation could be observed) or to a 1:1 ratio. For NMR titration experiments, various protein concentrations were used: ^15^N-labeled Hrp48-RRM1, RRM2, and RRM12 were titrated with different purchased RNA oligos, protein–protein interactions were tested by titrating ^15^N-labeled Hrp48 RRM12 with unlabeled Sxl dRBD3, Unr CSD123, CSD456, and CSD789 (details described in Table 2). CcpNmr Analysis, Sparky and scripts used in our lab were used to trace chemical shift perturbations and determine dissociation constants (*K*_D_) for interactions in the fast exchange regime. Individual *K*_D_ values of peaks were averaged to calculate global *K*_D_ values (Williamson, 2013). Errors were calculated from the individual fitting errors by error propagation. In the intermediate and slow exchange regime, the data fitting is erroneous and for comparative reasons the qualitative appearance of the spectra was assessed. Chemical shift perturbations were calculated according to

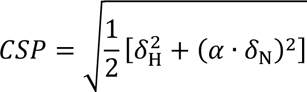

where 𝛼 = 0.2 was used based on the ratio between peak dispersion in the ^1^H and ^15^N dimensions (Williamson, 2013).

Measurements of longitudinal, *R*_1_ and transverse, *R*_2_ relaxation rates experiments for Hrp48-RRM12, Hrp48-RRM12-10-mer RNA complex and Hrp48-RRM12-9-mer RNA complex were acquired at proton Larmor frequencies of 700 MHz at 298 K using standard pulse sequences (Kay et al., 1989; Lakomek et al., 2012). The sample concentration was 80 µM for RRM12 in the apo state and 200 μM for RRM12 in the RNA bound state. The 9-mer and 10-mer RNA were added at 1.2-fold excess. For RRM12 alone, only 2 points were measured because of the aggregation tendency of RRM12 without RNA at elevated concentrations. For the *R*_1_ experiment, relaxation delays were 80 and 1200 ms and for the *R*_2_ experiment 16 and 80 ms. For RNA-bound Hrp48-RRM12 each experiment was recorded with 6 and 9 relaxation delays. For *R*_1,_ delays of 80, 400, 800, 1200, 1600, and 2000 ms were used with 80, 800, 1200 and 2000 ms as duplicates. For *R_2_* delays, 16, 32, 48, 64, 80, 96, 112, 128 and 144 ms relaxation delays were used with 32 ms as duplicates. For peak integration, error estimation and exponential fitting NMRViewJ was used (Johnson, 2004). Calculations of the relaxation rates from two-point measurements were completed in spreadsheets and errors were estimated through calculation of the standard deviation of the mean value of τ_c_ of each residue within a single RRM domain. The mean value was than calculated again for residues with τ_c_ values within the standard deviation. Residues outside of this range were considered as outliers. The experimental rotational correlation times were calculated according to (Fushman et al., 1994).

### Crystallization, data collection, and structure determination

Crystallization trials were performed using 3-lens 288-well crystallization plates using the sitting drop method, in which two sample concentrations/conditions can be tested with the same crystallization buffer. For this, 5-6 different 96-condition commercial screens were used at 4°C and at 20°C. Each well contained 0.1 μl sample and 0.1 μl crystallization condition solution, set up with a Mosquito LCP liquid handling robot. Several conditions yielded crystals for Hrp48-RRM1. For the final crystal that gave the highest resolution in X-ray diffraction, RRM1 of 5 mg/ml concentration in the 200 mM NaCl, 30 mM NaPi, 2 mM DTT, pH 6.5. buffer was mixed in a 1:1 ratio (100 nl:100 nl) with 0.2 M K_2_SO_4_ and 20% (w/v) PEG-3350 reservoir solution at 4°C. The sitting drop vapor diffusion method was used, and rod-shaped crystals started to nucleate overnight and kept growing further for 7-10 days. Crystals were soaked in mother liquor supplemented with 30% glycerol as a cryoprotectant prior to freezing.

Diffraction datasets were recorded at the P13 beamline at the German Electron Synchrotron (DESY), Hamburg, Germany. The crystals diffracted up to 1.15Å resolution.

The structure of RRM1 was solved by *ab initio* molecular replacement with the human hnRNP A1 RRM1 structure (pdb 1HA1, 40% sequence identity) as model using Phenix Phaser-MR (Adams et al., 2010). The initial model was built using Phenix AutoBuild (Terwilliger et al., 2008) and manual adjustments were executed with Coot (Emsley et al., 2010). The structure was further improved in iterative rounds of manual correction with Coot and restrained refinement with phenix.refine (Afonine et al., 2012). The crystal structure is deposited at the Protein Data Bank under the following accession code: 9EN7

### Molecular dynamics simulation of the RRM1-6-mer-3 complex

To simulate the spontaneous binding process (Krepl et al., 2022) of the 5′-UUAAG-3′ motif of the 6-mer-3 RNA (AUUAAG) to the RRM1 domain, we have constructed a system with the RNA positioned ∼20 Å away from the protein. Note that the 5′-terminal A_1_ of the full 6-mer-3 was skipped to increase sampling efficiency of the simulations. The initial coordinates of the protein and RNA were obtained from the X-ray structure of the RRM1 domain (PDB: 9EN7) and by NAB (Nucleic Acid Builder), respectively (Case et al., 2023). The OL3 (Zgarbová et al., 2011) and ff12SB (Maier et al., 2015) force fields were used to describe the RNA and protein, respectively. We have applied the recently introduced stafix approach (using scaling factor of 0.5) to eliminate spurious intramolecular RNA self-interactions occurring with the standard AMBER RNA simulation force field (Krepl et al., 2022). The RNA and the protein were immersed in an octahedral box of SPC/E water molecules (Berendsen et al., 1987) with minimal distance of 14 Å between the solutes and the box border. We have added the K^+^ and Cl^-^ ions (Joung and Cheatham, 2008) to neutralize the systems and obtain an ion concentration of ∼0.15 M. Prior to the production simulations, the systems were minimized and equilibrated (Krepl et al., 2018). The production simulations were then performed in constant pressure ensemble. Monte-Carlo barostat and Langevin thermostat were used to control the pressure and temperature, respectively (Case et al., 2023). We have performed nine 10-μs-long independent MD simulations, with different trajectories obtained by utilizing random seed numbers. The resulting trajectories visualized the RNA at different stages of binding to RRM1. By careful visual analysis, we have subsequently manually selected a binding motif showing the RNA stably bound at a location close to all the protein residues which exhibited chemical shift perturbations in NMR experiments. Note that due to structural complexity of the system and multiple-pathway nature of the binding process (Krepl et al., 2022), it is not feasible to carry out the selection based on any single collective variable (CV), such as the interaction energy of the complex. The natural multi-dimensionality of interactions also precludes application of CV-based enhanced sampling simulation methods. Long-term stability of the chosen binding motif was subsequently verified in two independent 10-μs-long MD simulations. Finally, the structure was used as our working model for the putative RRM1/6-mer-3 protein-RNA complex structure.

### Size exclusion chromatography-multi angle light scattering (SEC-MALS)

For SEC-MALS experiments different columns were used: a Superdex 200 Increase 10/300 GL or a Superdex 200 Increase 5/150 GL gel-filtration column coupled to the MiniDAWN and Optilab MALS system from Wyatt Technology. For each complex the single components (proteins and RNA alone) were measured first followed by the complex. After mixing and incubation on ice, the samples were centrifuged for 10 mins at 15000 rpm and then 50 μl were loaded onto the Superdex 200 Increase 5/150 GL or 100 μl onto the Superdex 200 Increase 10/300 GL columns. The minimal concentration was 1.0 mg/ml for each sample. The experiments were performed at room temperature in SEC buffer filtered twice through 0.22 μm pore-size filter. Data analysis was performed using the Astra 7 software (Wyatt Technology).

### *In vitro* translation assays

Wild type construct BLEF, as previously described (Gebauer et al., 2003), is composed of 69 nt of *msl-2* 5′ UTR sequence including site B and 46 nt of the *msl-2* 3′ UTR including sites E and F. For introducing mutations in the 3’ UTR, primers were created by QuikChange Primer Design (https://www.agilent.com/) and used for restriction-free cloning to create changes in the Region 5 and Region 6. Sxl-dRBD4 (amino acids 122–301 of *Drosophila melanogaster* SXL) was expressed in *Escherichia coli* as an N-terminal GST-tagged fusion protein and purified as described (Grskovic et al., 2003). The protein was dialyzed against buffer D (20 mM HEPES pH 8.0, 20% glycerol,1 mM DTT, 0.01% NP-40, 0.2 mM EDTA). BLEF mRNA derivatives were synthesized using T3 RNA polymerase (Ambion) and contained a 5′ m^7^GpppG cap and a poly(A) tail of 73 residues. mRNAs were purified by phenol-chloroform extraction and G_50_ columns (GE Heathcare). All mRNAs used in the same experiment were synthesized and quantified in parallel, and the concentration and quality confirmed by separation in agarose gels. *In vitro* translation assays were performed as described (Gebauer et al., 1999). Briefly, 17 ng of *msl-2* Firefly reporter mRNA and 10 ng Renilla luciferase mRNA, used as an internal control, were incubated with increasing amounts of GST-dRBD4 in a final volume of 10 µl. The reaction contained 40% *Drosophila* embryo extract, 60 µM amino acids, 16.8 mM creatine phosphate, 80 ng*/*µl creatine kinase, 24 mM HEPES pH7.5, 0.6 mM Mg(OAc)_2_ and 80 mM KOAc. The reaction was incubated at 25°C for 90 min, and the Firefly and Renilla luciferase activities were measured using the Dual Luciferase kit (Promega).

### Electrophoretic mobility shift assays

The GG-30-mer RNA oligo used for EMSAs was 3′ end-labeled with pCp-Cy-5 (Cyanine 5). For this, the following reaction mixture was combined in 20 µl final volume: 100 pmol RNA, 200 pmol pCp-Cy-5, 2 µl T4 RNA ligase (10x, 20 U), 2 µl DMSO, 1 mM ATP, 1 mM DTT, 10 mM MgCl_2_, 50 mM Tris-HCl (pH 7.5). The reaction was incubated at 16 °C overnight. Subsequently the reaction was purified by NaOAc/EtOH precipitation. To the reaction mixture, 4 µl (0.2 × volume) of 3 M sodium-acetate, 1.5 µl of GlycoBlue Coprecipitant (Invitrogen), and 59 µl (2.5 × volume) of ethanol cooled to −20°C was added and mixed by vortexing. The RNA was precipitated overnight at −70 °C, centrifuged at 4 °C and 13000 rpm. for 30 min, washed two times with −20°C 70% ethanol and once with 100% ethanol. The final amount and the labeling efficiency were measured by spectrophotometry using the NanoDrop^TM^ OneC. Electrophoretic mobility shift assays (EMSAs) were used to determine RNA-binding affinities in a semi-quantitative mode (Garner and Revzin, 1981). Recombinantly purified Hrp48-RRM12 was mixed with 20 nM Cy-5 labeled GG-30-mer probe in a buffer consisting of 50 mM Tris-HCl (pH 8.4), 100 mM NaCl, 1 mM EDTA, 250 mg/ml BSA, 10% glycerol, 0.05% Triton X-100 and 1 mM DTT in 12.5 µL reactions and incubated on ice for 30 min, protected from light. The concentration of proteins is indicated in the figure. The RNA – protein complexes were resolved on a 6% native 1 × TBE polyacrylamide gel for 35 min at 200 V. The gels were imaged at a Typhoon Trio Imager 9000 (GE Healthcare).

## Acknowledgements

We thank the local contacts at DESY Hamburg PETRA-3 (P12 beamline) for support. J.Š. and M.K. acknowledge support of the Czech Science Foundation (grant number 23-05639S). F.G. acknowledges support from the Spanish Ministry of Science and Innovation (MCIN) (grant PID2021-127948NB-I00 funded by MCIN / AEI / 10.13039/501100011033 / FEDER, UE) and the Catalan Agency for Research and Universities (SGR-Cat-2021-01215). J.H. gratefully acknowledges support from the Deutsche Forschungsgemeinschaft (DFG, HE 7291/5-1).

## Author contributions

Conceptualization: A.L., J.M., J.H.; Methodology: A.L., J.M., T.G., M.K., K.L., K.S., B.S., J.S., F.G., J.H.; Investigation: A.L., J.M., T.G., M.K., K.L., C.H., K.S., B.S.; Formal analysis: A.L., J.M., T.G., M.K., K.L., C.H., K.S., B.S., J.H.; Resources: J.S., F.G., J.H.; Writing-original draft: A.L., J.M., J.H.; Writing-review and editing: A.L., J.M., T.G., M.K., K.L., C.H., K.S., B.S., J.S., F.G., J.H.; Supervision: J.S., F.G., J.H.; Visualization: A.L., J.M., J.H.; Funding acquisition: J.H.

