## Supplementary Material for "The *Drosophila* RNA binding protein Hrp48 binds a specific RNA sequence of the *msl-2* mRNA 3’ UTR to regulate translation"

* These authors contributed equally

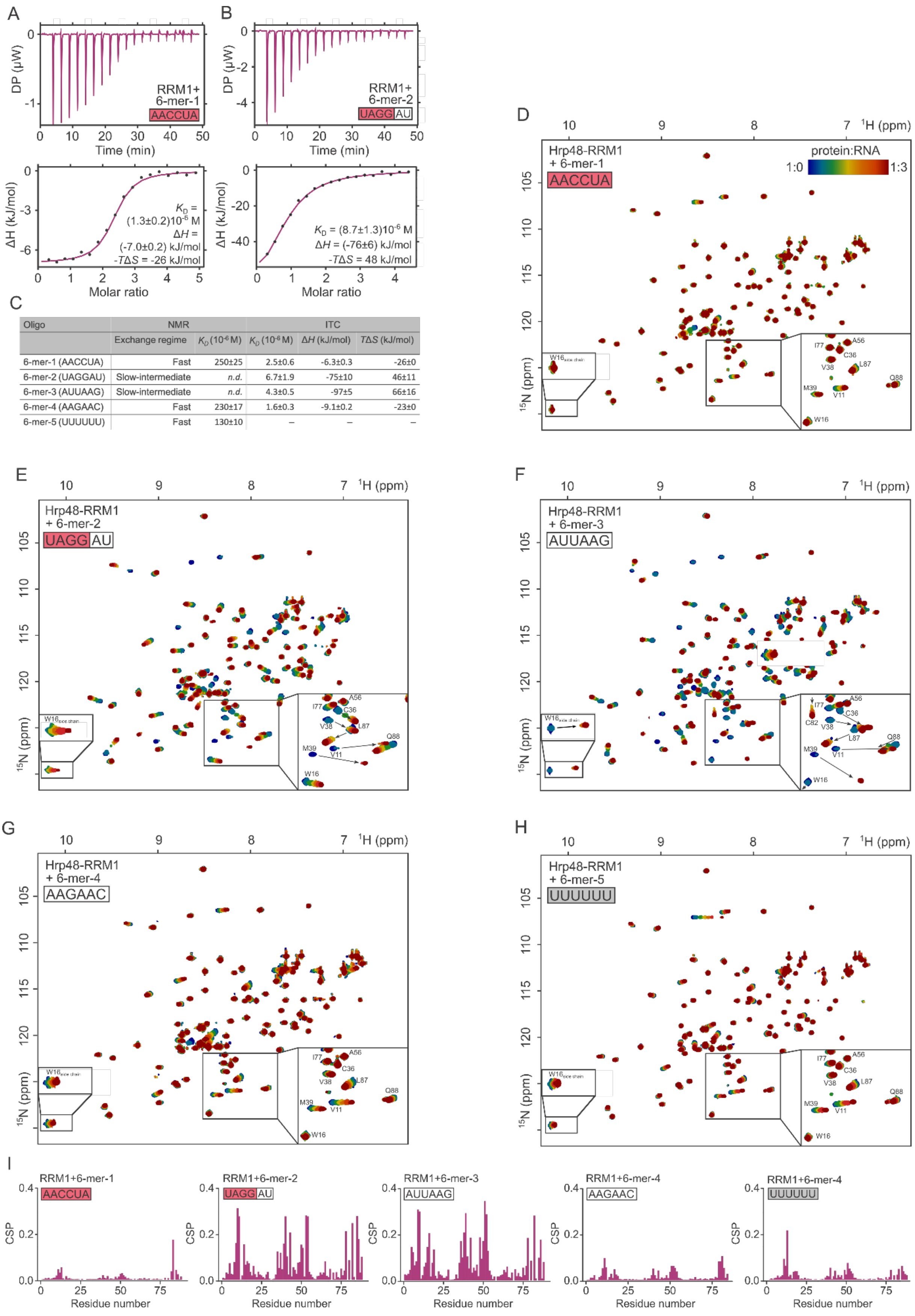


| Oligo | NMR | | ITC | | |
|---|---|---|---|---|---|
| | Exchange regime | $K_D$ ($10^{-6}$ M) | $K_D$ ($10^{-6}$ M) | $\Delta H$ (kJ/mol) | $T\Delta S$ (kJ/mol) |
| 6-mer-1 (AACCUA) | Fast | 250±25 | 2.5±0.6 | -6.3±0.3 | -26±0 |
| 6-mer-2 (UAGGAU) | Slow-intermediate | *n.d.* | 6.7±1.9 | -75±10 | 46±11 |
| 6-mer-3 (AUUAAG) | Slow-intermediate | *n.d.* | 4.3±0.5 | -97±5 | 66±16 |
| 6-mer-4 (AAGAAC) | Fast | 230±17 | 1.6±0.3 | -9.1±0.2 | -23±0 |
| 6-mer-5 (UUUUUU) | Fast | 130±10 | – | – | – |

**Figure S1:** Binding studies of Hrp48-RRM1. **A:** Isothermal titration calorimetry of 6-mer-1 titrated into RRM1. Data fitting yielded a dissociation constant of 1.3 μM, with modest heat release and positive entropy change. This is similar to the binding mode of RRM1 to 6-mer-4 (Figure 1). **B:** Isothermal titration calorimetry data of RRM1 and 6-mer-2, $K_D$ = 8.7 μM. The binding of this RNA involves a significant amount of heat release and negative entropy change, similarly to 6-mer-3 and RRM1. **C:** Table of NMR and ITC results of interaction studies between RRM1 and the 6-mers. The determination of the $K_D$ by fitting the NMR data in the slow-intermediate exchange regime is not possible, but the qualitative differences in the spectra clearly select these 6-mers as the strongest binding partners. Averaged ITC data were derived from replicate experiments, all data are shown on

Table *3*. Errors were calculated with error propagation except for $T\Delta S$, where the standard deviation of the averaged values is shown, because of the lack of experimental error. While the $K_D$ values determined by ITC are similar, the thermodynamic parameters suggest different binding modes between 6-mers. The titration of RRM1 with 6-mer-2 and 6-mer-3 happens with significant heat release compensated by a negative change in entropy, while titration with 6-mer-1 and 6-mer-4 induce a small heat release and a positive change in entropy. 6-mer-5 binding was not strong enough to allow reliable fitting of ITC data. **D-H:** Complete $^{15}N$,$^{1}H$-HSQC titration spectra for the Hrp48-RRM1 titration experiments with 6-mers, zoomed into the same region as in Figure 1 and into the tryptophan (W16) side chain signal. The coloring scheme indicated in D is the same for each titration. **H:** 6-mer-5 is a control of which the sequence is not related to the other 6-mers. RRM1 binds this motif, albeit with weak affinity. **I:** CSP-plots of 6-mer NMR titration experiments.

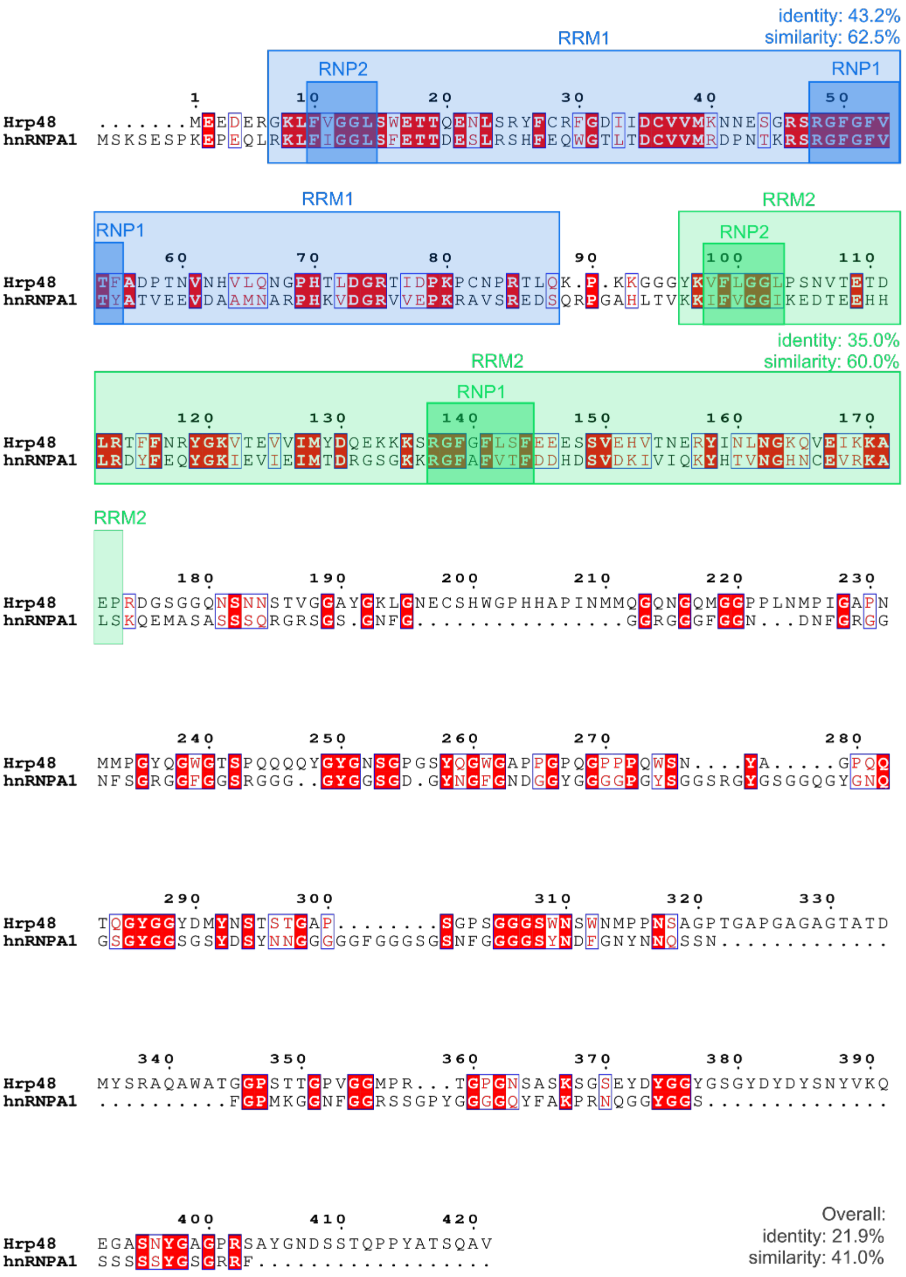


**Figure S2:** Sequence alignment of Hrp48 and hnRNP A1. The identical sequences are highlighted in red; the similar amino acids are indicated with red fonts. The RRM domains are marked by blue (RRM1) and green (RRM2) boxes, and the conserved RNP-sequences with darker blue and green respectively. The sequence similarity and identity was calculated with Emboss Needle (Madeira et al., 2022). The figure was prepared using Clustal Omega (Madeira et al., 2022) multiple sequence alignment tool and ESPript (Robert and Gouet, 2014).

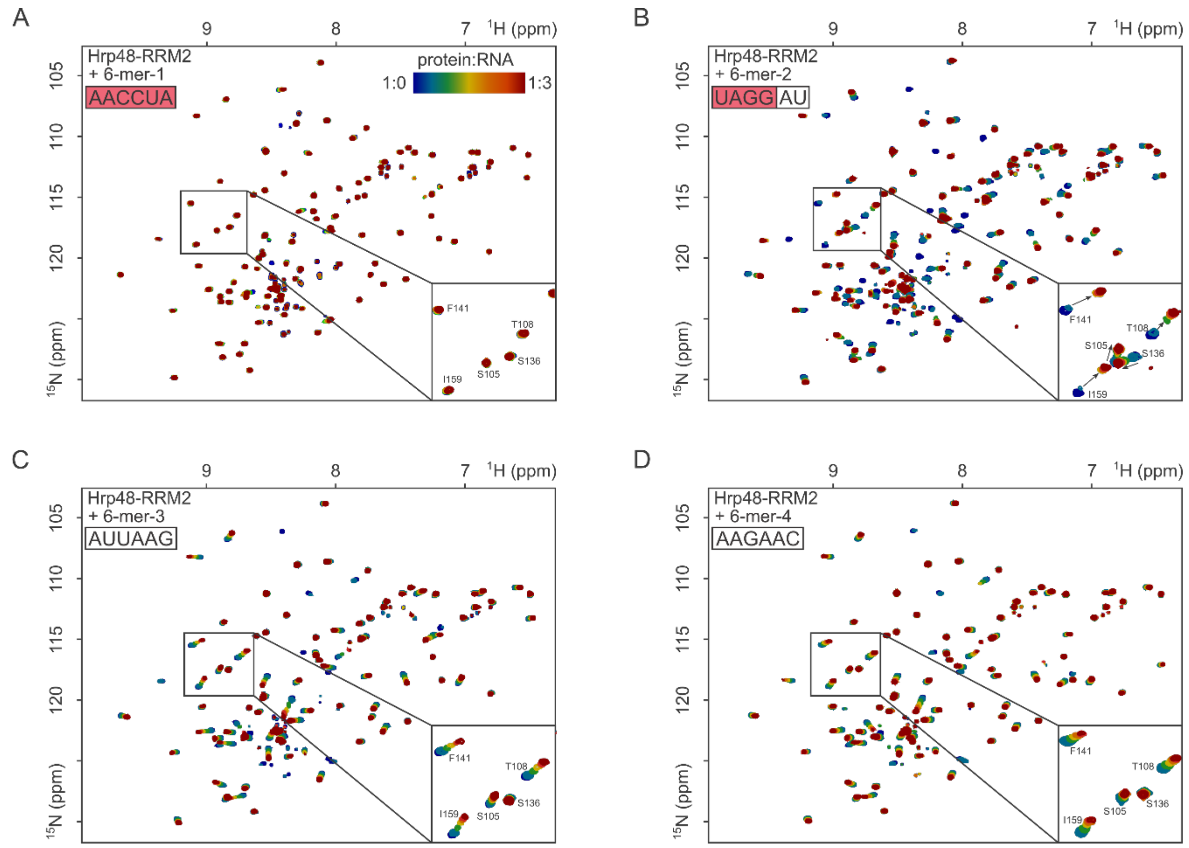


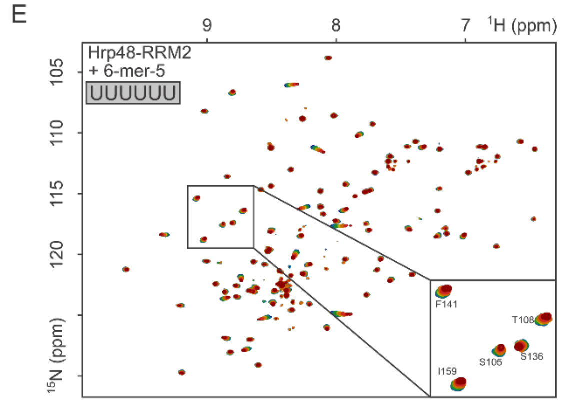


F

| Oligo | NMR | |
|---|---|---|
| | Exchange regime | $K_D$ ($10^{-6}$ M) |
| 6-mer-1 (AACCUA) | Fast | 597±132 |
| 6-mer-2 (UAGGAU) | Slow-intermediate | *n.d.* |
| 6-mer-3 (AUUAAG) | Intermediate | 22.6±2.9 |
| 6-mer-4 (AAGAAC) | Intermediate | 20.7±1.9 |
| 6-mer-5 (UUUUUU) | Fast | 317±40 |

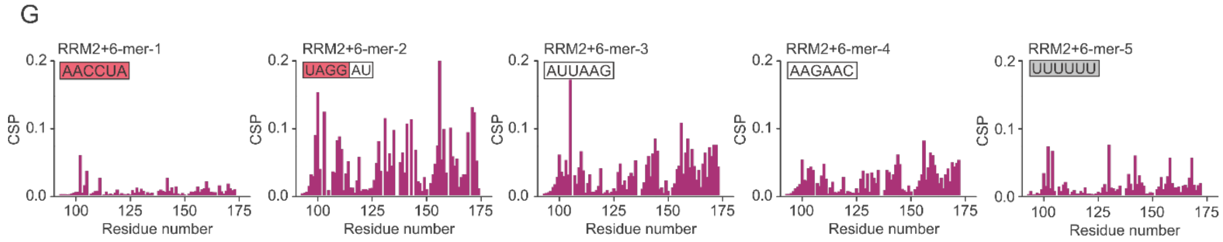


**Figure S3:** Binding studies of Hrp48-RRM2. **A-E:** Complete $^{15}$N,$^{1}$H-HSQC titration spectra for Hrp48-RRM2 interaction experiments with the different 6-mer RNAs, with a zoom-in into the same region as shown in Figure 2. The coloring scheme indicated in A is the same for each titration. **F:** Table of NMR results of interaction studies between RRM2 and the 6-mers. Based on the appearance of the spectra, 6-mer-2 is the strongest binding partner for RRM2. **G:** CSP plots of 6-mer NMR titration experiments.

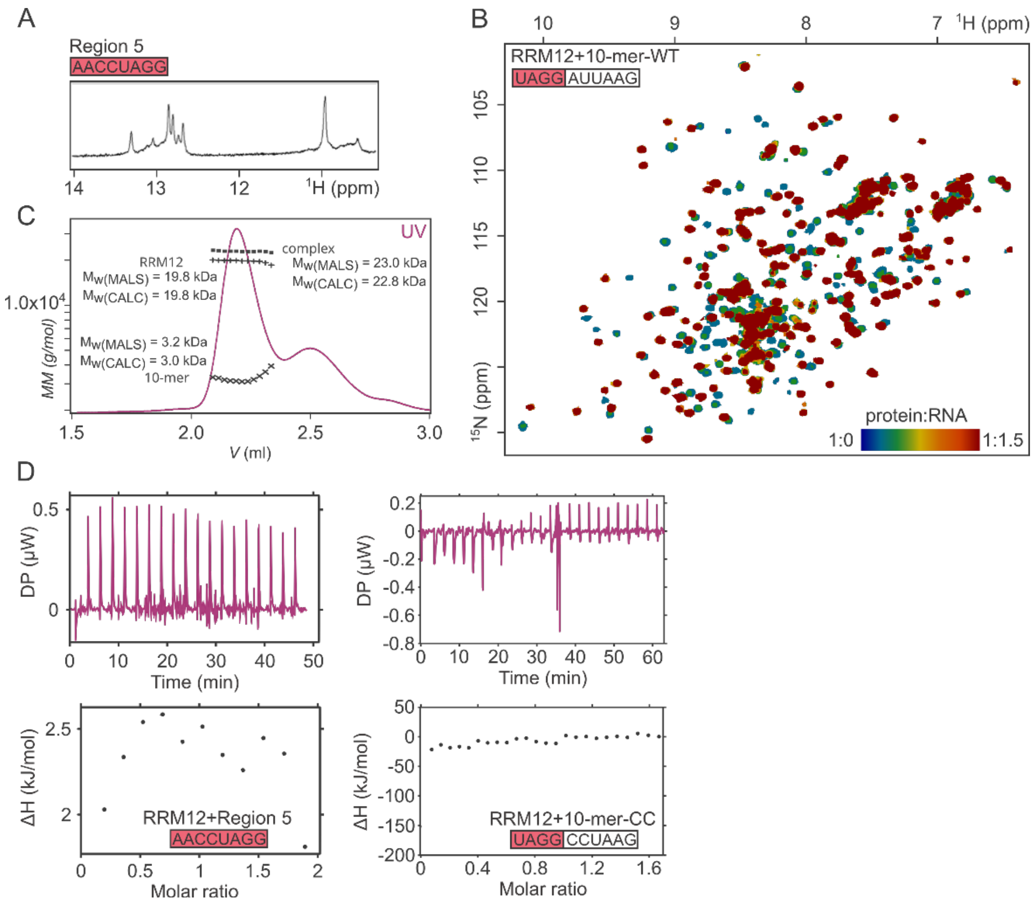


**Figure S4:** Binding studies of Hrp48- RRM12. **A:** We confirmed duplex formation of Region 5 by observing imino–hydrogen peaks in a one-dimensional $^{1}$H–NMR spectrum, which is strong evidence of self-association. **B:** $^{15}$N,$^{1}$H-HSQC titration spectra for Hrp48-RRM12 interaction experiments with the 10-mer-WT with peaks exhibiting chemical shift perturbation in the slow exchange regime, indicating sub micromolar binding affinity. **C:** SEC–MALS UV chromatogram and conjugate analysis of an injection of Hrp48-RRM12 + 1.2 eq 10-mer-WT. The 10-mer-WT and RRM12 forms a 1:1 complex. **D:** Left panel: ITC of Hrp48-RRM12 and Region 5 RNA construct which was suggested previously as the *msl-2* binding site of Hrp48. Although it would be expected that a tandem RRM would bind with nanomolar affinity, we could not confirm a strong interaction by ITC or NMR. Right panel: ITC of 10-mer-CC mutant RNA and Hrp48-RRM12. The binding affinity decreased significantly thus reliable fitting was not feasible. Measuring with higher protein concentration was not possible due to oligomerization tendency of Hrp48.

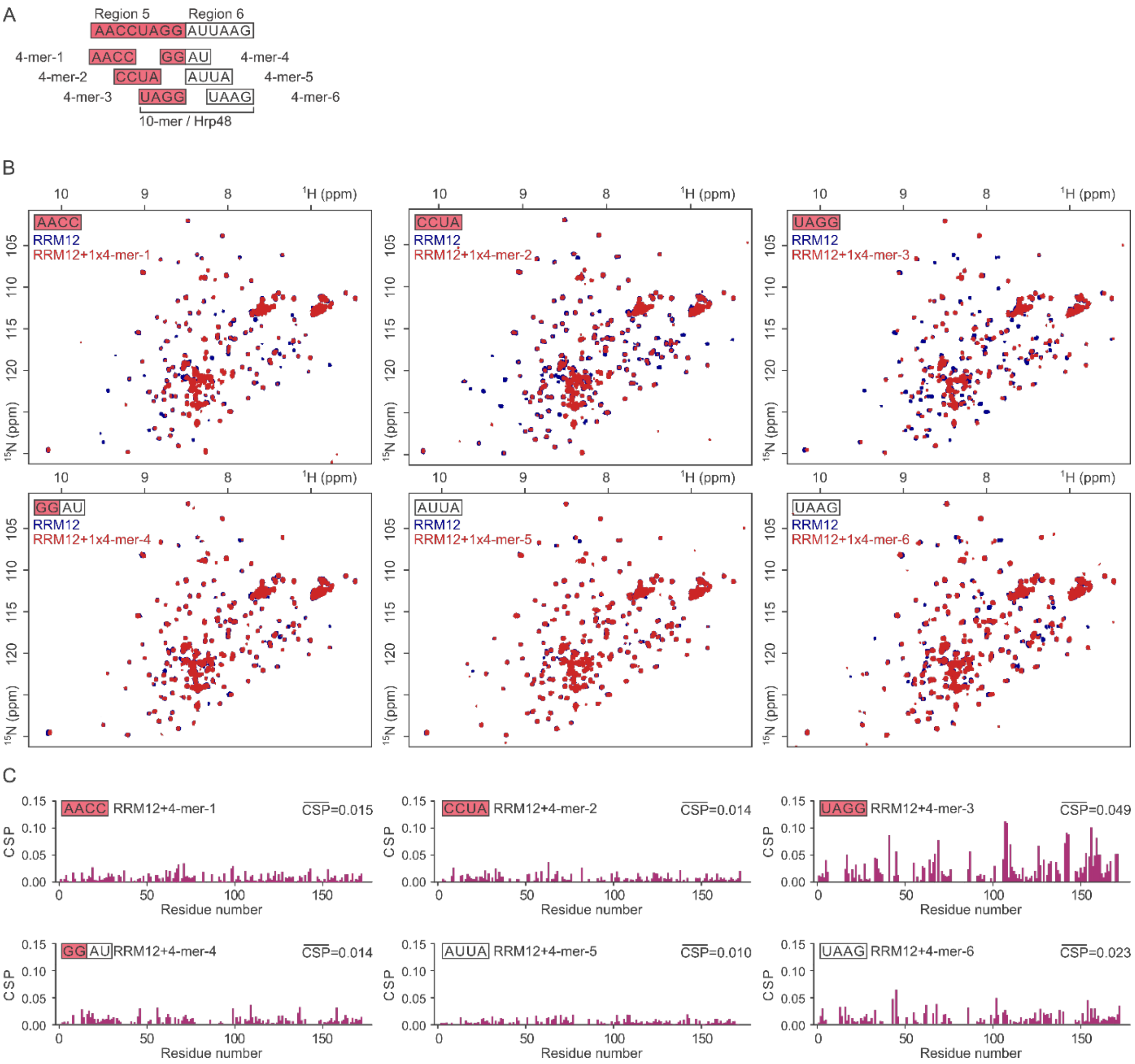


**Figure S5:** Refinement of the RNA motif bound by Hrp48-RRM12. **A:** 4-mers covering both Region 5 and 10-mer used in NMR titrations. **B:** $^{15}N$,$^{1}H$-HSQC NMR spectra of free RRM12 (blue) and RRM12 with one equivalent of respective 4-mer (red). At 1:1 ratio the RNA binding of 4-mer-3 exhibits the strongest chemical shift perturbations, followed by 4-mer-6. **C:** The chemical shift perturbations of RRM12 residues upon addition of one molar equivalent 4-mer RNA. The average CSP was calculated from the CSP values greater than the standard deviation of all CSP values. This was calculated for each experiment separately.

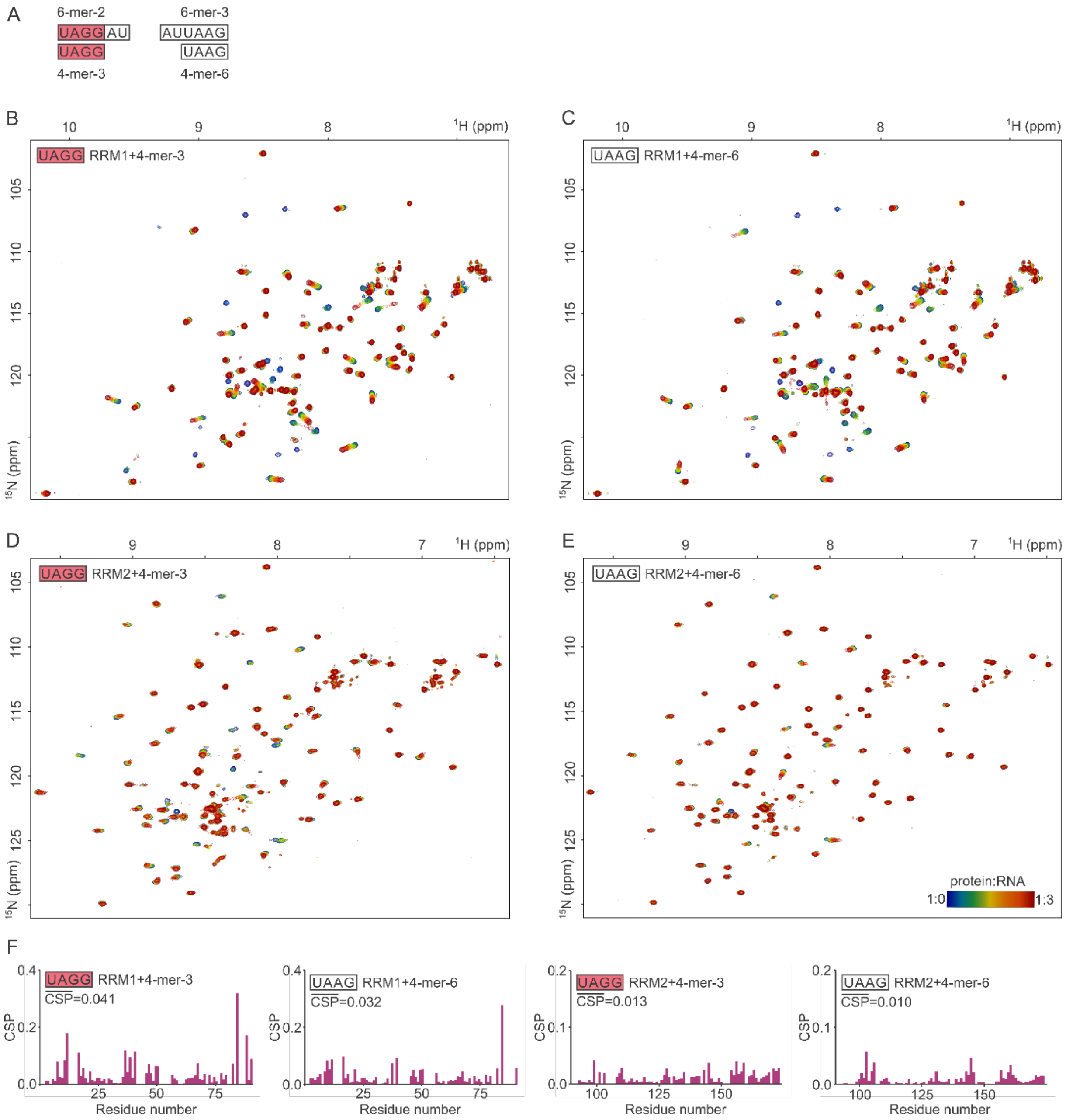


**Figure S6:** NMR titration of RRM single domains using shortened 4-mer RNA constructs. **A:** Sequences of 4-mers derived from 6-mer-2 and 6-mer-3. **B-C:** $^{15}$N,$^{1}$H-HSQC NMR spectra of Hrp48-RRM1 NMR titration experiments titrated with 4-mers. RNA was added to three-fold excess. **D-E:** $^{15}$N,$^{1}$H-HSQC NMR spectra of Hrp48-RRM2 NMR titration experiments titrated with 4-mers. RNA was added to three-fold excess. The color bar indicates the amount of RNA added during the experiments and is valid for all NMR spectra B-E. **F:** CSP plots of NMR titration experiments of B-E.

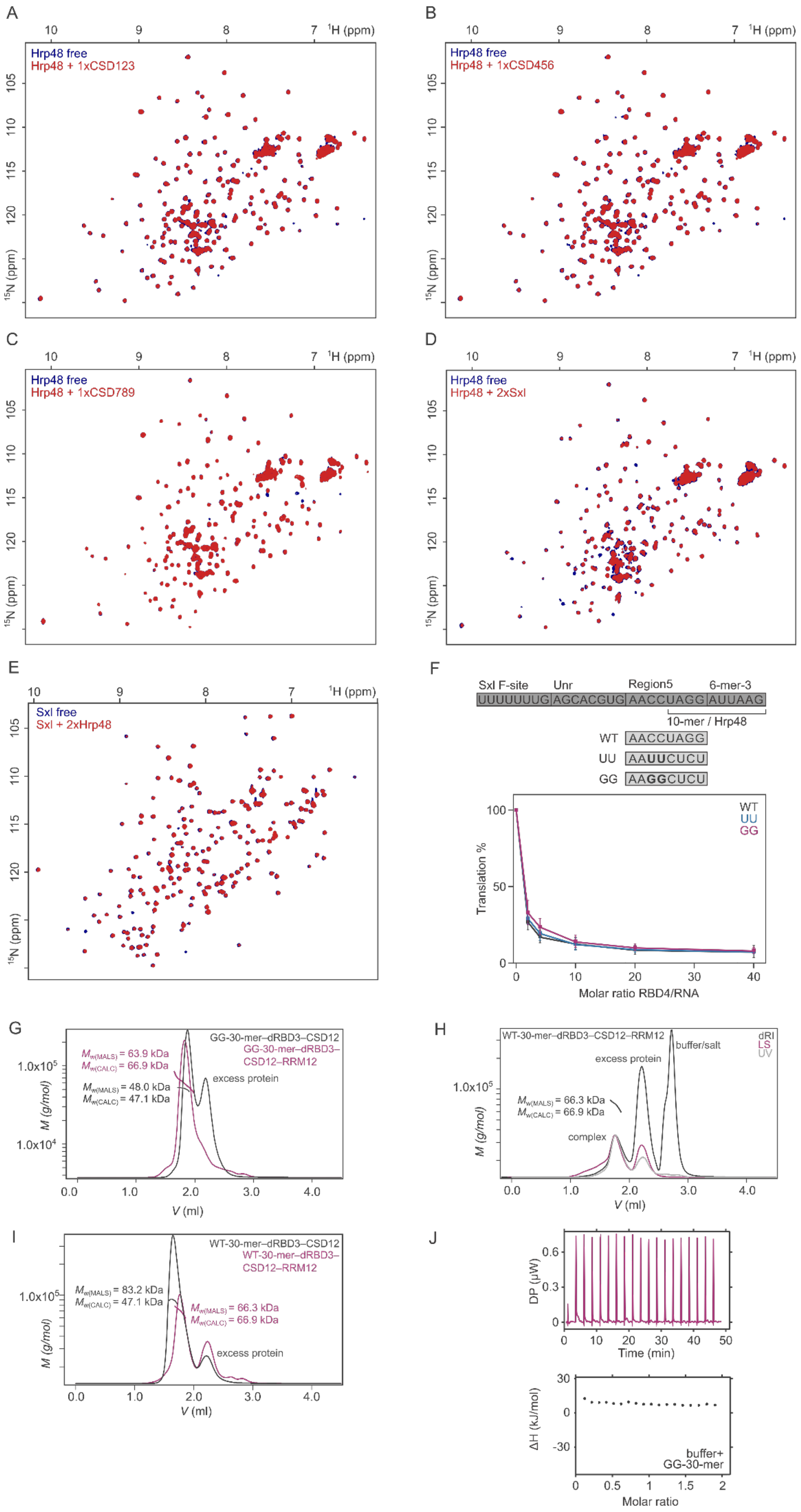

A
Hrp48 free
Hrp48 + 1xCSD123
B
Hrp48 free
Hrp48 + 1xCSD456
C
Hrp48 free
Hrp48 + 1xCSD789
D
Hrp48 free
Hrp48 + 2xSxl
E
Sxl free
Sxl + 2xHrp48
$^{1}$H (ppm)
$^{15}$N (ppm)
F
Sxl F-site
Unr
Region5
6-mer-3
UUUUUUUG AGCACGUG AACCUAGG AUUAAG
10-mer / Hrp48
WT AACCUAGG
UU AAUUCUCU
GG AAGGCUCU
WT
UU
GG
Translation %
Molar ratio RBD4/RNA
G
GG-30-mer–dRBD3–CSD12
GG-30-mer–dRBD3–CSD12–RRM12
$M_{w(MALS)}$ = 63.9 kDa
$M_{w(CALC)}$ = 66.9 kDa
$M_{w(MALS)}$ = 48.0 kDa
$M_{w(CALC)}$ = 47.1 kDa
excess protein
M (g/mol)
V (ml)
H
WT-30-mer–dRBD3–CSD12–RRM12
dRI
LS
UV
buffer/salt
excess protein
$M_{w(MALS)}$ = 66.3 kDa
$M_{w(CALC)}$ = 66.9 kDa
complex
M (g/mol)
V (ml)
I
WT-30-mer–dRBD3–CSD12
WT-30-mer–dRBD3–CSD12–RRM12
$M_{w(MALS)}$ = 83.2 kDa
$M_{w(CALC)}$ = 47.1 kDa
$M_{w(MALS)}$ = 66.3 kDa
$M_{w(CALC)}$ = 66.9 kDa
excess protein
M (g/mol)
V (ml)
J
DP (μW)
Time (min)
ΔH (kJ/mol)
buffer+
GG-30-mer
Molar ratio

**Figure S7:** Complex formation of Hrp48-RRM12, Sxl-dRBD3, Unr-CSD12 and 30-mer RNA. **A-E:** $^{15}N$,$^{1}H$-HSQC NMR spectra of $^{15}N$ labeled Hrp48-RRM12. Blue: free protein, red: free protein and an unlabeled Hrp48, Sxl or Unr construct. No significant chemical shift perturbations are observable upon addition of Sxl, Hrp48 or Unr to Hrp48 or Sxl. The weak line broadening effect for some of the peaks is possibly due to unspecific interactions. This corroborates that Hrp48-RRM12 does not interact with structured domains of Sxl and Unr in the absence of RNA. **F:** Mutation of C3 and C4 of Region 5 does not impair the translation repression activity. *In vitro* translation assays were performed for two constructs modified at C3 and C4 to UU or GG, represented on the graph by bold fonts. The assays were performed with increasing amounts of recombinant Sxl dRBD4. Co-translated Renilla luciferase was used as an internal control and as a reference for normalization of the reporter Firefly luciferase signal. The data was plotted as relative percentage normalized to the experiment without Sxl addition. The standard deviations of three replicate experiments are represented by the error bars. **G-I:** SEC–MALS UV chromatograms of the different complexes of *msl-2*, Hrp48, Sxl and Unr. The markers indicate the molecular weights of the molecules/complexes in the peaks. **G:** SEC–MALS UV chromatograms of GG-30-mer–dRBD3–CSD12 (grey, ratio: 1:2:3) and GG-30-mer–dRBD3–CSD12–RRM12 (purple, ratio: 1:2:3:2) complexes using proteins in excess. **H:** SEC–MALS chromatograms of the formation of the GG-30-mer–dRBD3–CSD12–RRM12 complex. Purple: light scattering (LS), dark grey: differential refractive index (dRI), light grey: UV absorption. **I:** SEC–MALS UV chromatograms of WT-30-mer–dRBD3–CSD12 (grey, ratio: 1:2:3) and WT-30-mer–dRBD3–CSD12–RRM12 (purple, ratio: 1:2:3:2) complexes using proteins in excess. The WT-30mer in complex with dRBD3 and CSD12 forms a dimer through the palindromic region of the WT-30-mer. This opens up when RRM12 is added and a 1:1:1:1 complex forms. **J:** Control experiment: 30-mer RNA titrated into buffer. The peaks are equal in magnitude to each other and similar to the peaks at the end of the curves on Figure 5E. This indicates no effect of dilution on the RNA which would complicate the evaluation of the ITC data on Figure 5E

**Table 1:** Data collection and refinement statistics.

| | Hrp48_RRM1 |
|---|---|
| **Wavelength** | |
| **Resolution range** | 31.82 – 1.191 (1.233 – 1.191) |
| **Space group** | P 21 21 21 |
| **Unit cell** | 38.344 45.4996 57.026 90 90 90 |
| **Total reflections** | 416881 (41304) |
| **Unique reflections** | 32496 (856) |
| **Multiplicity** | 12.8 (13.1) |
| **Completeness (%)** | 91.18 (26.81) |
| **Mean I/sigma(I)** | 18.30 (1.27) |
| **Wilson B-factor** | 14.59 |
| **R-merge** | 0.06499 (1.536) |
| **R-meas** | 0.06773 (1.597) |
| **R-pim** | 0.01881 (0.4347) |
| **CC1/2** | 1 (0.609) |
| **CC*** | 1 (0.87) |
| **Reflections used in refinement** | 29733 (857) |
| **Reflections used for R-free** | 2000 (58) |
| **R-work** | 0.1627 (0.2193) |
| **R-free** | 0.1617 (0.2151) |
| **CC(work)** | 0.972 (0.855) |
| **CC(free)** | 0.976 (0.840) |
| **Number of non-hydrogen atoms** | 803 |
| **macromolecules** | 670 |
| **ligands** | 28 |
| **solvent** | 121 |
| **Protein residues** | 84 |
| **RMS(bonds)** | 0.006 |
| **RMS(angles)** | 0.88 |
| **Ramachandran favored (%)** | 98.78 |
| **Ramachandran allowed (%)** | 1.22 |
| **Ramachandran outliers (%)** | 0.00 |
| **Rotamer outliers (%)** | 0.00 |
| **Clashscore** | 2.98 |
| **Average B-factor** | 24.44 |
| **macromolecules** | 21.65 |
| **solvent** | 66.07 |

Statistics for the highest-resolution shell are shown in parentheses.

**Table 2:** Relative and absolute concentrations used for NMR titrations.

| Analyte protein in NMR tube | Conc. NMR tube (µM) | Titrants | Molar ratios |
|---|---|---|---|
| Hrp48-RRM1 | 20/50/100 | 6-mer-1, 6-mer-2, 6-mer-3, 6-mer-4, 6-mer-5, 4-mer-3, 4-mer-6 | 1.0:0.0 – 1.0:2.5 or 1.0:3.0 |
| Hrp48-RRM2 | 20/50 | 6-mer-1, 6-mer-2, 6-mer-3, 6-mer-4, 6-mer-5, 4-mer-3, 4-mer-6 | 1.0:0.0 – 1.0:2.5 or 1.0:3.0 |
| Hrp48-RRM12 | 100,<br><br>50 | 10-mer, Sxl-dRBD4, Unr-CSD123, Unr-CSD456, Unr-CSD789,<br>4-mers | 1.0:0.0 – 1.0:2.5<br><br>1.0:0.0 – 1.0:1.0 or 1.0:1.5 |
| Sxl-dRBD3 | 100 | Hrp48-RRM12 | 1.0:0.0 – 1.0:2.5 |

Table 3 Detailed parameters and results of ITC measurements.

| Exp. name | Replicates | Conc. syringe (µM) | Conc. cell (µM) | *N* (sites) | *N* (error) | *K*d (µM) | *K*d (µM, error) | Δ*H* (kJ/mol) | −*T*Δ*S* (kJ/mol) | Injection no. | Injection vol. (µl) | Shown in figure |
|---|---|---|---|---|---|---|---|---|---|---|---|---|
| Hrp48-RRM1 + 6-mer-1 | 2 | 6-mer-1 766–800 | RRM1 30.0 | 2.4<br>3.1 | $45\cdot10^{-3}$<br>$121\cdot10^{-3}$ | 1.5<br>3.5 | 0.4<br>1.2 | -7.2±0.3<br>-5.3±0.4 | -25.6<br>-25.4 | 19 | 2.0 | S1 |
| Hrp48-RRM1 + 6-mer-2 | 3 | 6-mer-2 189–550 | RRM1 17.5–20.0 | 0.8<br>0.8<br>0.9 | $35\cdot10^{-3}$<br>$76\cdot10^{-3}$<br>$94\cdot10^{-3}$ | 8.7<br>4.0<br>7.4 | 1.3<br>2.2<br>2.9 | -76±6<br>-65±12<br>-85±16 | 48<br>35<br>56 | 19/25 | 2.0/1.5 | S1 |
| Hrp48-RRM1 + 6-mer-3 | 4 | 6-mer-3 235–601 | RRM1 23.0–25.0 | 0.8<br>0.6<br>0.4<br>0.7 | $17\cdot10^{-3}$<br>$18\cdot10^{-3}$<br>$13\cdot10^{-3}$<br>$17\cdot10^{-3}$ | 5.0<br>6.1<br>2.6<br>3.4 | 0.6<br>0.5<br>0.4<br>0.4 | -105±5<br>-100±5<br>-107±6<br>-74±3 | 75<br>71<br>76<br>43 | 19/25 | 2.0/1.5 | 1 |
| Hrp48-RRM1 + 6-mer-4 | 3 | 6-mer-4 650–1400 | RRM1 17.5–30.0 | 1.7<br>1.6<br>2.4 | $32\cdot10^{-3}$<br>$12\cdot10^{-3}$<br>$37\cdot10^{-3}$ | 1.0,<br>0.8,<br>3.0 | 0.3,<br>0.1,<br>0.5 | -9.7±0.3<br>-10.4±0.1<br>-7.3±0.2 | -24.0<br>-23.9<br>-23.7 | 19/25 | 2.0/1.5 | 1 |
| Hrp48-RRM12 + Region 5 | 1 | Region 5 1000 | RRM12 60.0 | No binding detectable by ITC. | | | | | | 19 | 2.0 | S4 |
| Hrp48-RRM12 + 10-mer-WT | 2 | 10-mer 171 | RRM12 22.0 | 0.6<br>0.6 | $2.2\cdot10^{-3}$<br>$1.21\cdot10^{-3}$ | 0.02<br>0.02 | $2.9\cdot10^{-3}$<br>$1.6\cdot10^{-3}$ | -275±2.6<br>-282±1.5 | 232<br>238 | 25 | 1.5 | 3 |
| Hrp48-RRM12 + 9-mer | 2 | 9-mer 182 | RRM12 21.6 | 0.7<br>0.7 | $3.2\cdot10^{-3}$<br>$2.5\cdot10^{-3}$ | 0.02<br>0.01 | $3.7\cdot10^{-3}$<br>$3.2\cdot10^{-3}$ | -235±0.3<br>-226±0.7 | 191<br>181 | 25 | 1.5 | 3 |
| Hrp48-RRM12 + 10-mer-CC | 2 | 10-mer-CC 190 | RRM12 21.6 | No binding detectable by ITC. | | | | | | 25 | 1.5 | S4 |
| Hrp48-RRM12 + GG-30-mer | 3 | GG-30-mer | RRM12 | Complex binding mode of multiple binding sites. | | | | | | 19/25 | 2.0/1.5 | 5 |
| Hrp48-RRM12 + GG-30-mer–Sxl-dRBD3–Unr-CSD12 complex | 4 | complex 283–300 | RRM12 19.0–29.3 | 0.5<br>0.6<br>0.5 | $18\cdot10^{-3}$<br>$14\cdot10^{-3}$<br>$9.7\cdot10^{-3}$ | 0.9<br>1.8<br>1.9 | 0.2<br>0.3<br>0.3 | -124±7<br>-126±6<br>-120±4 | 90<br>94<br>128 | 19/25 | 2.0/1.5 | 5 |
